# Intertwined trade-offs coordinate *Drosophila* midgut mitosis vs. endoreplication and host defense vs. dysplasia

**DOI:** 10.1101/615104

**Authors:** Vasilia Tamamouna, Myrofora Panagi, Andria Theophanous, Maria Demosthenous, Maria Michail, Markella Papadopoulou, Savvas Teloni, Chrysoula Pitsouli, Yiorgos Apidianakis

## Abstract

Inflammatory signaling supports host defense against infection, not only through immune cells, but also via regeneration of damaged tissue. Heightened regeneration, nevertheless, predisposes for all types of cancer and thus a trade-off exists between regeneration capacity and long-term tissue homeostasis. Here, we study the role of tissue-intrinsic regenerative inflammatory signaling in stem cell mitosis of the adult *Drosophila* midgut at the baseline and the infected state and its impact on intestinal host defense to infection and stem cell-mediated dysplasia. Through a quantitative genetics screen we find that stem cell mitosis is positively linked with the expression of *eiger, Delta, upd3* and *vein* in the midgut, as well as with dysplasia and host defense, but negatively with enterocyte endoreplication. We provide evidence that intertwined trade-offs fine-tune midgut homeostasis, according to which stem cell mitosis through *cyclin E* in stem cells promotes the optimal host defense to infection, unless dysplasia ensues. However, *cyclin E* in enteroblasts promotes enterocyte endoreplication and counterbalances stem cell mitosis and dysplasia, providing an alternative but less efficient mechanism to support host defense.

## Introduction

Although a link between inflammatory microenvironment and tumor initiation and progression is clearly established, the mechanisms underlying cell interactions in the tumor and its microenvironment remain unclear. Known germline mutations account only for a small percentage of cancers developing with age, whereas the vast majority of cancer-promoting factors are due to spontaneous somatic mutations facilitated by inflammation (Hanahan, 2011). Inflammation of the colonic mucosa is a key predisposing factor for developing colon cancer and is usually accompanied by high mitosis and DNA damage (Balkwill, 2001; Lasry, 2016). Inflammation is linked directly through STAT signaling and indirectly through reactive oxygen species to tissue regeneration and mutation (Taniguchi, 2015; Karin, 2016; Panayidou, 2013). Strikingly, the level of tissue-intrinsic mitosis per se is a key factor in cancinogenesis in humans (Tomasetti, 2015; Tomasetti, 2107). Similarly, proliferative activity in the aging *Drosophila* midgut correlates positively with dysplasia, but also with longevity, with maximal lifespan when intestinal proliferation is reduced, but not completely inhibited (Biteau, 2010). Nevertheless, the genetic basis of the tissue-intrinsic quantitative control on mitosis remains obscure (Wu, 2016). Tissue homeostasis in higher eukaryotes is established via coordinated events of cell proliferation, death and differentiation. The *Drosophila* midgut, like its mammalian counterpart, is frequently challenged by a plethora of abiotic and biotic stresses, which can damage the intestinal epithelial barrier leading to pathogenesis. Thus, homeostatic mechanisms must be tightly regulated in terms of epithelial cell turnover and shedding of damaged epithelial cells (Buchon, 2013; Paterson, 2014; Karin, 2016). The *Drosophila* midgut is maintained by pluripotent intestinal stem cells (ISCs) that self-renew and give rise to transient enteroblasts (EBs), which will terminally differentiate into polyploid enterocytes (ECs) (Micchelli, 2006; Ohlstein, 2006). ISCs also give rise to the pre-enteroendocrine cell (pre-EEs) population, the EE progenitors (Zeng, 2015). *Drosophila* midgut ISC mitosis is controlled by key signaling pathway ligands, including *Delta, upd1*-3, *vein, dilp3*,6 and *eiger* (Doupe, 2018), but mitosis can be diverted towards dysplasia due to deregulation of many of the same signaling pathways these ligands may control (Apidianakis 2009; Biteau 2008). Adult midgut dysplasia in *Drosophila* has been characterized by the widespread, irreversible and progressive loss of proper cell differentiation due to accumulation of groups of ISC-like and EE cells (Apidianakis, 2009; Biteau, 2008; Resende, 2018), and it is to be contrasted with the rare ISC-like/EE large tumors caused by spontaneous loss of heterozygosity of *notch* in old flies (Siudeja, 2015).

A common property of differentiating *Drosophila* cells that need to cope with tissue development and homeostasis is endoreplicative cell growth. Endoreplication or endocycling is an evolutionary conserved biological process during which cells undergo many cycles of DNA replication without dividing (Kluza, 2011; Tamori, 2014; Shu, 2018). A prerequisite for the transition from mitotic cycles to endocycles is the inhibition of mitosis, whereby Cyclin E (CycE) oscillations regulated by Cyclin dependent kinase 1 (Cdk1) control G1/S transition in mitotic cells and G/S transition in endoreplicating cells. CycE facilitates the expression of many S-phase control genes (Shu, 2018; Duronio, 1994) and its levels fluctuate between the G1, S, G2 and M phases of the cell cycle, so that CycE accumulates at S phase and is absent during the G2 phase to allow the formation of the pre-endoreplication complex required for the next round of DNA synthesis (Weiss, 1998). Thus, cells undergo endoreplication depending on the expression levels of CycE. Importantly, malfunctions in CycE regulation induce chromosomal instability and facilitate development of many cancers such as bone, lung, intestine, brain, liver and breast (Bortner, 1997; Donnella, 1999; Malumbres, 2001).

Endoreplication, in addition to inefficient cell mitosis, contributes significantly to tissue regeneration upon damage. The JAK/STAT and the EGFR pathways are activated in both the *Drosophila* midgut stem cells to induce their mitosis and in EBs and young ECs to promote endoreplication (Jiang, 2009; Xiang, 2017). In the damaged adult epidermis, the Hippo pathway induces compensatory polyploidization, and in the hindgut, pylorus endoreplication increases in response to apoptosis (Losick, 2013). Similarly, the Insulin/IGF-like pathway in necessary for compensatory endoreplication in response to follicular epithelium cell loss (Tamori, 2013). The same pathway is necessary for midgut growth upon pupal eclosion and upon feeding on rich vs. poor media (O’Brien, 2011; Choi, 2011).

In this study, we investigated how cell growth and proliferation are coordinated in the *Drosophila* midgut epithelium at a given homeostatic state, either at baseline or during an established infection. We found that intestinal mitosis varies greatly as a function of the fly strain’s genetic background and certain tissue morphological characteristics may adapt to the high versus low mitosis status. Our quantitative genetics study demonstrates that epithelial intestinal mitosis sustained via Notch, JAK-STAT, EGFR and TNFR pathway ligands improve host defense to infection, but predispose for dysplasia, which compromises host defense. Nevertheless, we noticed an inverse correlation between stem cell mitosis and endoreplication, which is controlled by cycE in both types of cells, so that midgut progenitors undergo endoreplication to compensate for cell loss, when mitotic activity is limited and *vice versa*.

## Results

### Isogenized *Drosophila* strains exhibit extreme differences in ISC mitosis before and upon infection

To genetically dissect the phenotypic variation of intestinal mitosis, we screened 153 sequenced wild-type strains from the *Drosophila* Genetics Reference Panel (DGRP) (McKay, 2012) measuring the number of mitotic cells per midgut upon oral infection with the human opportunistic pathogen *Pseudomonas aeruginosa* (Apidianakis, 2009). Z-score analysis of the mitotic index (Fig. 1a) underscored a great variation in mitosis, of up to 5 standard deviations of the mean, among the strains tested. Upon repeated examination of strains exhibiting extreme mitosis, we selected 22 of them: 11 that were consistently highly mitotic, and 11 that were lowly mitotic (Fig. 1b). These are color highlighted in Fig. 1a and listed in Fig 1f. To pinpoint causal factors for the differences between the two groups of strains in terms of intestinal inflammation and dysplasia, we undertook a multi-parametric approach assessing fly-associated bacteria, defense to infection, endoreplication, dysplasia and tissue-intrinsic inflammatory signaling.

**Figure 1.**
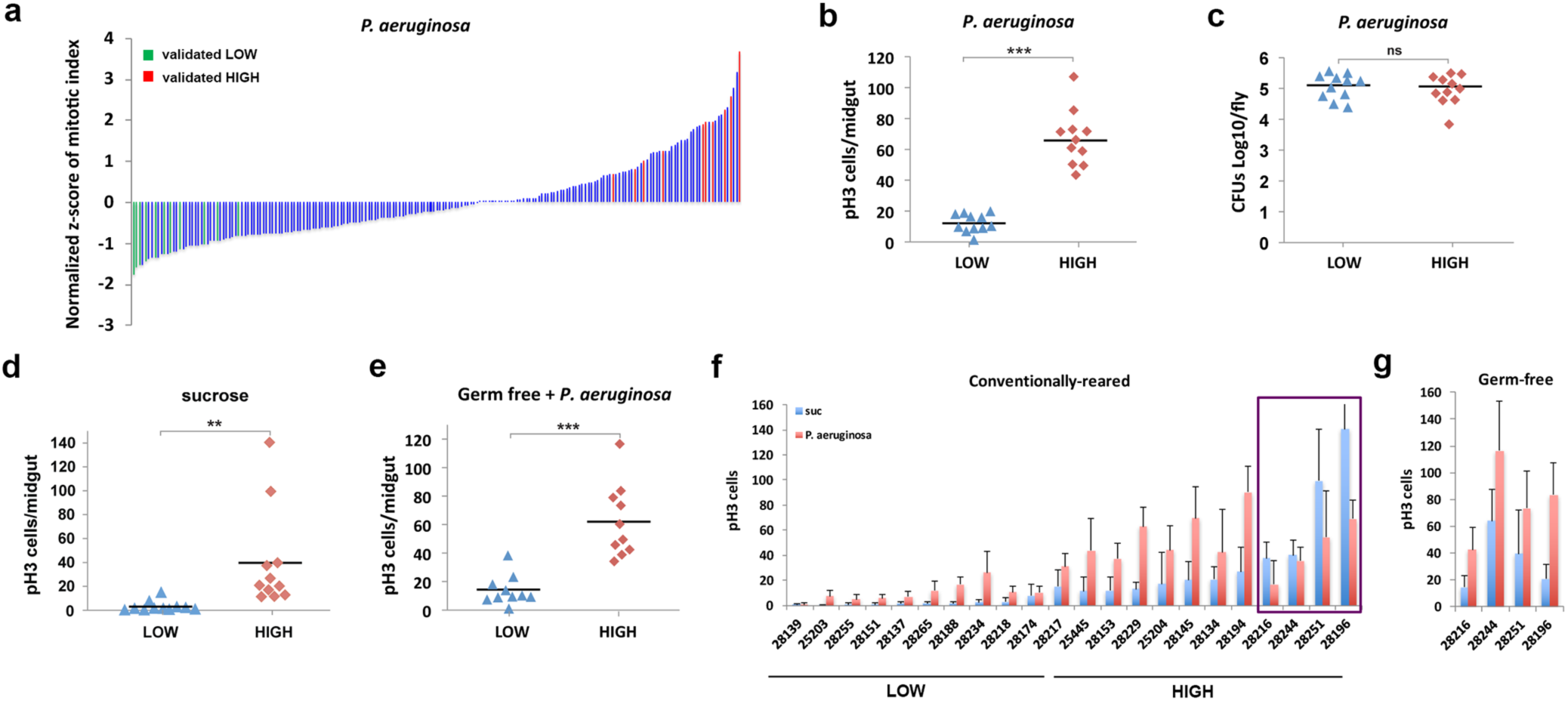
Phenotypic variation among 153 DGRP lines pinpoints extreme mitotic strains. **a.** Z-score of the 153 DGRP lines exhibiting variation of >5 times the standard variation. Green and red bars indicate validated low and high mitosis strains, respectively. **b.** Extreme “low” (28139, 25203, 28255, 28151, 28137, 28265, 28188, 28234, 28218, 18174, 28217) and “high” (25445, 28153, 28229, 25204, 28145, 28134, 28194, 28216, 28244, 28251, 28196) mitosis strains exhibit differential midgut mitosis (pH3 cells/midgut) upon *P. aeruginosa* infection. **c.** Colony forming units (CFUs) of *P. aeruginosa* for each of the 22 extreme strains grouped as “low” and “high”. **d.** Low and high mitosis strains exhibit differential midgut mitosis without infection. **e.** Germ-free low and high mitosis strains exhibit differential midgut mitosis upon infection. **f-g.** Mitosis per midgut of conventionally-reared extreme strains without (sucrose only) and upon infection with *P. aeruginosa*. The 4 strains in rectangle appear to lack induction by infection, when conventionally-reared (f), but they are inducible, when grown as germ-free (g). Significance in b-e is assessed by t-test: ns *p*>0.05; * 0.01<*p*≤ 0.05; ** 0.001<*p*≤ 0.01; *** *p*≤ 0.001.

### Highly mitotic strains are inherently prone to ISC mitosis and more resilient to infection

To assess if higher mitosis supports host defense to infection we assessed high and low mitotic strains in two different ways: (a) either individually for each of the 22 homogeneous groups of flies or (b) from a single group of flies of mixed genotype by sampling 5 flies for each of the 22 strains (highly mitotic expressing GFP, while lowly mitotic did not). In both experiments we noticed that highly mitotic strains cope significantly better with lethal intestinal infection (Suppl. Fig. 1). Nevertheless, the two extreme groups of strains contain comparable numbers of pathogenic bacteria upon infection (Fig. 1c), indicating that resilience to *P. aeruginosa* infection (the process of better tolerating pathogenic microbes without eliminating them) rather than resistance (the process of better inhibiting pathogenic microbe growth in the host) is taking place (Schneider, 2008; Ferrandon, 2013).

Mitosis is inducible by microbiota and pathogenic bacteria in both the high and low mitosis strains. Nevertheless, midgut mitosis of the two groups of strains is significantly different, not only after, but also before infection (Fig. 1d), and microbiota is not a key factor for the difference in mitosis, because germ-free fly strains retain their difference in mitosis between the two groups and upon infection (Fig. 1e). The effect of microbiota is evident though in 4 of the highly mitotic fly strains, in which mitosis is inducible upon *P. aeruginosa* infection when germ-free, but not when conventionally-reared, because microbiota accounted for high level of mitosis (Fig. 1f-g). Nevertheless, the effect of microbiota on mitosis of these 4 strains does not change their classification as highly mitotic (1f-g). Thus, mitosis is inducible by microbiota and pathogenic bacteria in both the high and low mitosis strains. Nevertheless, even in the absence of microbial stimulus, the baseline mitosis level differs between the two groups of strains.

### Highly and lowly mitotic strains respond similarly to infection in terms of midgut size, cell damage and shedding

To assess if gut dimensions are affected by the differential level of mitosis in the highly vs. lowly mitosis strains, we assessed differences before and upon infection in midgut size. We noticed that, despite initial differences, the response of the two groups of strains to infection is similar. That is, midguts of the highly mitotic strains are generally longer (Suppl. Fig. 2a-c), but, similarly to lowly mitotic ones, they do not increase their length upon infection (Suppl. Fig. 2d-e). In addition, highly mitotic strains have posterior midguts that are larger in width (Suppl. Fig. 2f-g). Nevertheless, highly and lowly strains respond similarly to infection by increasing their anterior and posterior midgut width to comparable levels (Suppl. Fig. 2h-i). Thus, baseline differences in size rather than a differential tissue growth upon infection distinguishes the highly vs. lowly mitotic strains.

**Figure 2.**
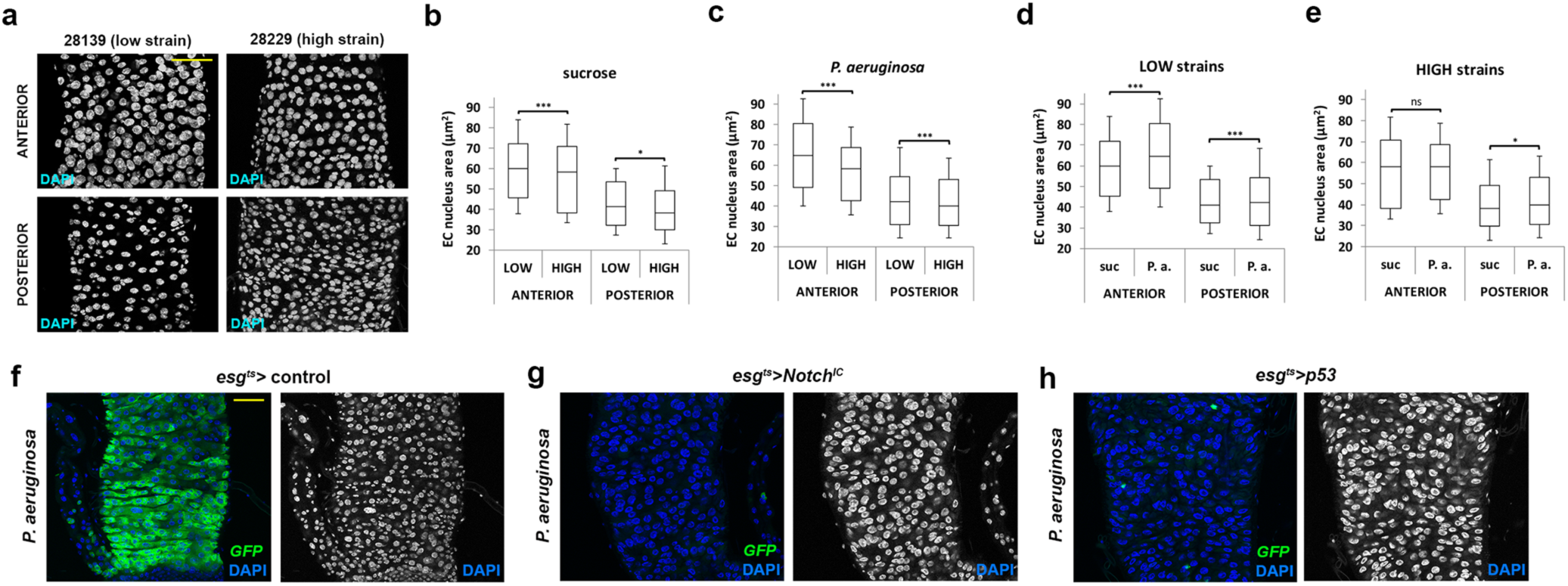
Intestinal ISC mitosis inversely correlates with EC endoreplication. **a.** Representative confocal images of anterior and posterior midguts of a low (28139) and a high (28229) mitosis strain in baseline conditions. DAPI stains the cell nuclei. **b-c.** Quantification of anterior and posterior EC nucleus size of midguts displaying low (b) and high (c) mitosis in baseline conditions and upon *P. aeruginosa* infection. **d-e.** Comparison of EC nucleus size between uninfected (d) and infected (e) conditions in low and high mitosis strains. Significance in b-e is assessed by t-test: ns *p*>0.05; * 0.01<*p*≤ 0.05; ** 0.001<*p*≤ 0.01; *** *p*≤ 0.001. **f-h.** Elimination of ISCs leads to increased EC size: compare the nucleus size of infected control midguts (f) to progenitor-specific expression of *Notch*^*IC*^ that promotes differentiation of ISCs (g) and *p53* that kills ISCs (h). Green: *esg-Gal4 UAS-GFP tub-Gal80*^*ts*^; Blue: DAPI. Scale bar 50 um.

To assess if differences in mitosis are a result of differences in enterocyte (EC) damage and exfoliation, we used methylene blue to transiently stain the *Drosophila* midgut epithelium, allowing the measurement of the decoloration rate for each fly strain (Suppl. Fig. 3a-c). The decoloration rate increases upon *P. aeruginosa* infection (Suppl. Fig. 3d), which is known to kill midgut ECs by apoptosis (Apidianakis, 2009). Nevertheless, we did not observe any differences in the decoloration rates of the highly vs. the lowly mitotic strains upon infection (Suppl. Fig. 3e-f). In agreement to this, *puckered* (*puc*), a downstream target of the JNK pathway predominantly induced in stressed or damaged ECs (Apidianakis, 2009). is expressed at similar levels between the two groups of strains (Suppl. Fig. 3g, Table 1). We conclude that the level of EC damage and exfoliation upon infection does not distinguish the two extreme groups of fly strains.

**Table 1:** Summary of RT-qPCR gene expression experiments in extreme “low” and “hig h” mitosis strains. Significant expression differences are indicated in red. Comparisons between low vs high strains in baseline conditions (sucrose), low vs high strains upon infection (P. a.) and all extreme strains (low and high combined) in sucrose vs infected conditions are shown. Grey shading indicates candidate regulators identified by GWAS. The *p*-values are calculated using the Student’s t-test. Biological triplicates are used for all samples.

| Gene name | Symbol | $p$ baseline (suc)<br>Low vs High | $p$ infected (*P.a.*)<br>Low vs High | $p$ Low+High<br>suc vs *P.a.* |
| --- | --- | --- | --- | --- |
| *Attacin A* | *AttA* | 0.837 | 0.724 | **0.053** |
| *branchless* | *bnl* | 0.550 | 0.685 | 0.084 |
| *breathless* | *btl* | 0.291 | 0.538 | 0.480 |
| *Cyclin E* | *CycE* | 0.939 | 0.066 | 0.766 |
| *Death-associated inhibitor of apoptosis 1* | *Diap1* | 0.780 | 0.651 | 0.348 |
| *decapentaplegic* | *dpp* | 0.211 | 0.472 | 0.625 |
| *Delta* | *DI* | **0.030** | 0.870 | 0.415 |
| *Diptericin A* | *Dipt* | 0.187 | 0.577 | 0.059 |
| *Drosomycin-like 3* | *Dro3* | 0.707 | 0.461 | 0.483 |
| *Dual oxidase* | *Duox* | 0.459 | 0.721 | 0.837 |
| *Ecdysone-inducible gene 3* | *Impl3* | 0.982 | 0.806 | 0.508 |
| *eiger* | *egr* | **0.049** | 0.391 | 0.105 |
| *grindelwald* | *grnd* | 0.266 | 0.371 | 0.366 |
| *hedgehog* | *hh* | 0.076 | **0.003** | **0.003** |
| *Immune-regulated catalase* | *Irc* | **0.027** | 0.908 | 0.463 |
| *Insulin-like peptide 3* | *Dilp3* | 0.507 | 0.232 | 0.951 |
| *Insulin-like peptide 7* | *Dilp7* | 0.408 | 0.730 | 0.762 |
| *Keren* | *krn* | 0.537 | 0.810 | **0.007** |
| *NADPH oxidase* | *Nox* | 0.965 | 0.978 | 0.133 |
| *puckered* | *puc* | 0.821 | 0.180 | 0.150 |
| *spatzle* | *spz* | 0.447 | 0.700 | 0.221 |
| *spitz* | *spi* | 0.659 | 0.952 | 0.279 |
| *unpaired 3* | *upd3* | **0.014** | 0.854 | **0.000** |
| *vein* | *vn* | **0.001** | 0.243 | 0.245 |
| *wengen* | *wgn* | **0.050** | **0.036** | 0.143 |
| *wingless* | *wg* | 0.568 | 0.635 | 0.899 |
| *CG4991* | *CG4991* | **0.011** | **0.012** | 0.782 |
| *CG8475* | *CG8475* | 0.264 | 0.496 | 0.860 |
| *Fidgetin* | *Fign* | 0.088 | 0.071 | 0.759 |
| *slamdance* | *sda* | 0.597 | 0.552 | 0.154 |
| *Sno oncogene* | *Snoo* | 0.106 | 0.488 | 0.127 |
| *Sorting nexin 6* | *Snx6* | 0.363 | 0.095 | 0.793 |

**Figure 3.**
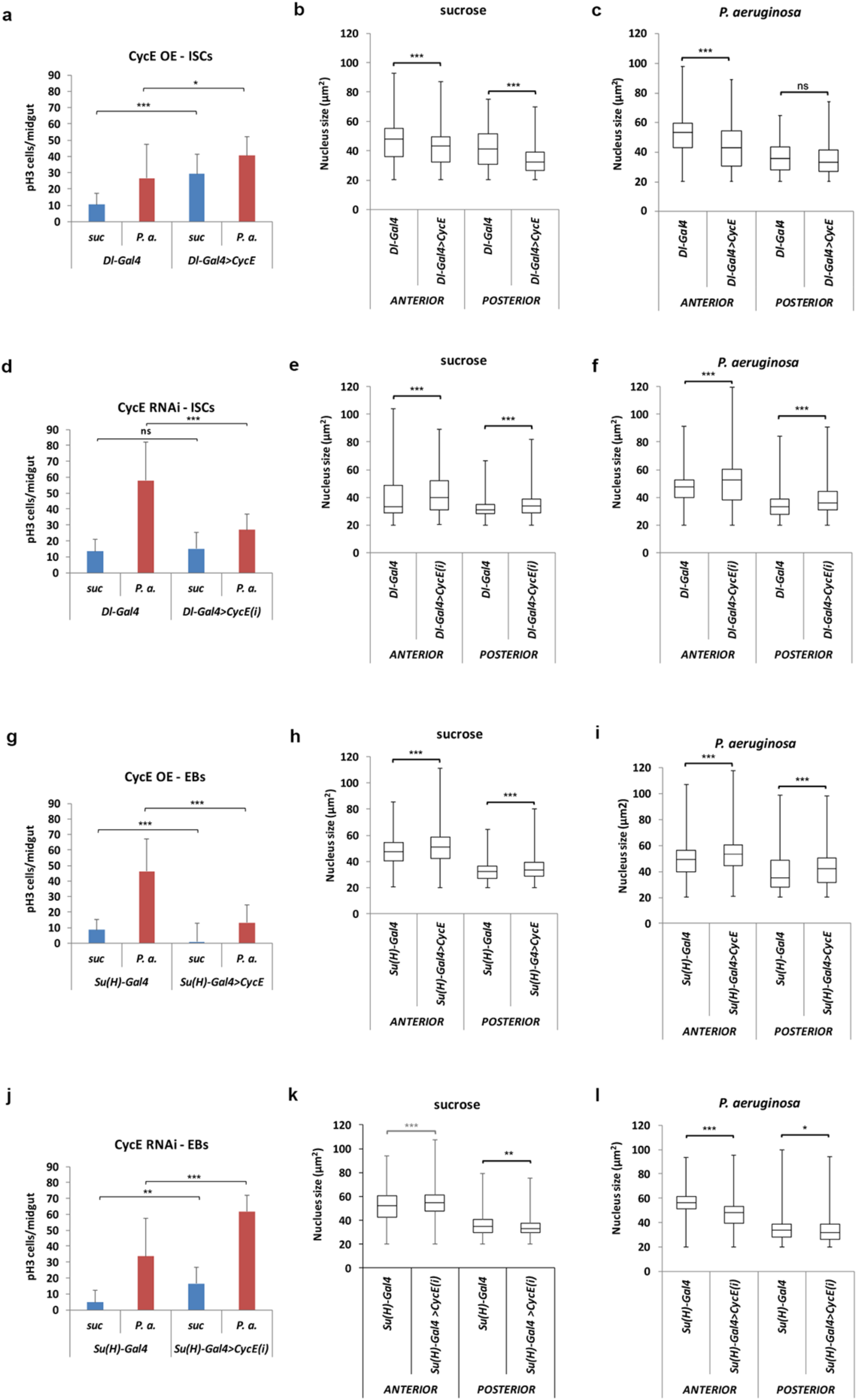
Increasing ISC mitosis induces EC endoreplication and vice versa. **a-c.** Midgut mitosis (a) and EC endoreplication in baseline (b) and infected (c) conditions upon *cycE* overexpression in ISCs using *Dl-Gal4*. **d-f.** Midgut mitosis (d) and EC endoreplication in baseline (e) and infected (f) conditions upon *cycE* RNAi knockdown in ISCs using *Dl-Gal4*. **g-h.** Midgut mitosis (g) and EC endoreplication in baseline (h) and infected (i) conditions upon *cycE* overexpression in EBs using *Su(H)-Gal4*. **j-l.** Midgut mitosis (j) and EC endoreplication in baseline (k) and infected (l) conditions upon *cycE* RNAi knockdown in EBs using *Su(H)-Gal4*. Significance is assessed by t-test: ns *p*>0.05; * 0.01<*p*≤ 0.05; ** 0.001<*p*≤ 0.01; *** *p*≤ 0.001.

### Inverse correlation between ISC mitosis and EC endoreplication at a given homeostatic demand

To address whether the comparable increase in midgut width of both lowly and highly mitotic strains upon infection can be attributed to counterbalance by endoreplication, we measured the nucleus size of the ECs in the anterior and posterior midgut of all 22 strains. EC specific measurements were obtained by setting a threshold of >20um^2^ in cell nucleus surface to exclude the diploid ISC, EB and EE populations (Supplementary Fig. 4). We found that endoreplicating ECs of lowly mitotic strains have bigger nuclei compared to highly mitotic strains, both in the anterior and posterior midgut, either at baseline or upon infection, indicating an inverse relationship between ISC mitosis and EC endoreplication at a given homeostatic demand (Fig. 2a-e). Thus, both groups of strains respond to infection by increasing their width at comparable levels (Suppl. Fig. 2h-i), but low mitosis strains are more prone in inducing endoreplication as a compensatory mechanism for tissue growth.

To further explore the proliferation-endoreplication interplay, we ablated genetically the ISC population in the adult midguts by expressing a constitutively active form of Notch protein lacking the extracellular domain (Notch^IC^) or by the overexpression of the pro-apoptotic protein p53 specifically in progenitor cells (Esg^+^) (Fig. 2f-h). While infection expands the progenitor pool in wild type flies, Esg+ cells were barely evident upon Notch^IC^ or p53 overexpression in adults. The absence of Esg+ cells resulted in substantially enlarged nuclei compared to the control sample, demonstrating that in the absence of mitotic cells infected ECs multiply their DNA as a compensatory mechanism to meet tissue homeostasis demands.

### CycE controls ISC mitosis and EC endoreplication cell autonomously and non-autonomously

CycE is a key regulator of cell cycle S phase in *Drosophila*. Downregulation of CycE levels in embryos inhibits DNA replication in both endoreplicative and mitotic cells (Knoblich, 1994), and overexpression of CycE in salivary glands induces DNA synthesis in endoreplicative cells when nutrients are low (Britton, 1998). Accordingly, we set out to assess if genetic manipulation of CycE levels in proliferating ISCs and differentiating EBs could affect their functions. We found that overexpression of CycE in ISCS via the *Dl-Gal4* increased mitotic activity by ∼2-fold, as indicated by pH3 staining, in the infected and non-infected epithelium, but reduced the nucleus size of ECs in the anterior and the posterior midgut at baseline before (Fig. 3a-c). Upon infection cycE in ISCs also reduced endoreplication in the anterior, but only tentatively so in the posterior, which we attribute to the excessive mitosis resulting in dysplasia (Fig. 4i). Consistently, downregulation of CycE specifically in the ISCs produced a ∼2-fold decrease in mitosis under infectious conditions and a concurrent increase in nucleus size of ECs in the anterior and the posterior midgut before and upon infection (Fig. 3d-f). On the other hand, CycE overexpression in transient EBs using the *Su(H)-Gal4* driver reduced intestinal mitosis by ≥4-fold upon infection, but increased EC nucleus size both in the anterior and the posterior midgut in the presence and absence of infection (Fig. 3g-i). Similarly, downregulation of CycE in EBs resulted in a ∼2-fold decrease in mitosis upon infection and an increase in endoreplication, with a notable exception of the uninfected anterior midgut (Fig. 3j-l). We conclude that CycE promotes mitosis cell autonomously in the ISCs and suppresses endoreplication non-cell autonomously, and *vice versa* CycE promotes endoreplication cell autonomously in EBs and suppresses mitosis non-autonomously.

**Figure 4.**
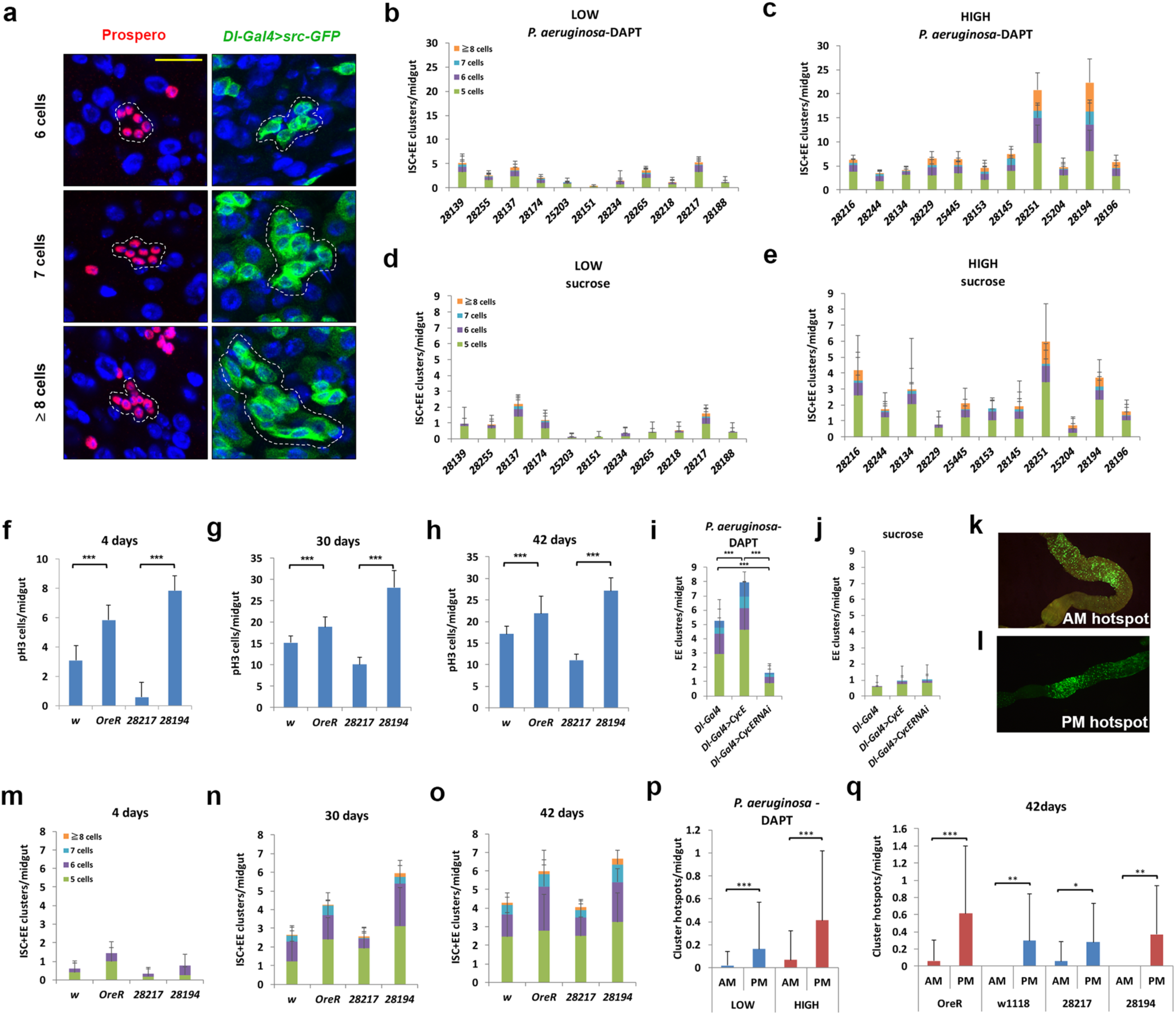
ISC and EE dysplastic cell clusters in chemically-treated and infected fly midguts, and in midguts upon aging. **a.** Prospero (red) and *Dl-Gal4 UAS-srcGFP* (green) cell clusters of 6, 7 and ≥8 cells. **b-c.** The 11 “High” vs. 11 “Low” mitosis strains differ in cluster numbers per female midgut in the presence of infection/DAPT. **d-e.** The 11 “High” vs. 11 “Low” mitosis strains differ in cluster numbers per female midgut in the absence of infection; both t-test assessed differences are significant (*p*<0.05). **f-h**, **m-o.** Differences in mitotic cells (f-h; t-test assessed) and ISC+EE clusters (m-o; chi-square tested) per female midgut during aging of 4 strains of different genetic background bearing the *Dl-Gal4 UAS-srcGFP* transgenes. **i-j.** *CycE* and *CycE*^*RNAi*^ expression in ISCs with *Dl-Gal4* in the presence (d) and absence (g) of infection/DAPT; all chi-square tested differences between control and transgene expression are significant at *p*<0.001. **k-l.** Hotspots of *Dl-Gal4 UAS-srcGFP*-positive cell clusters in the anterior and posterior midgut. **p-q.** Enumeration of cluster hotspots in the anterior and posterior midgut in DAPT/infected (p) and 42-day old flies (q); t-tested: ns *p*>0.05; * 0.01<*p*≤ 0.05; ** 0.001<*p*≤ 0.01; *** *p*≤ 0.001. Scale bar 25 um.

Next, we sought to assess CycE expression in EBs and ISCs along the midgut. Analysis of FlyGut-seq data (Buchon, 2013) showed a higher expression of CycE in the EBs vs. ISCs in the anterior midgut (an EB/ISC ratio of 2,88 for R1 and 1,56 for R2), but the opposite is noted in the posterior midgut (an EB/ISC ratio of 0,7 in R4 and 0,44 in R5) (Supplementary Fig. 5a). This may explain the bigger EC nuclei of the anterior compared to posterior midgut (Fig. 2 and Fig. 3) and the higher mitosis we noted in the posterior compared to the anterior midgut (Supplementary Fig. 5b-c). Taken together, these results suggest that the relatively higher CycE in the anterior EBs boosts their endoreplication, while the relatively higher CycE in the posterior ISCs boosts their proliferation.

### CycE-promoted endoreplication in EBs improves host defense to infection

To assess the impact of endoreplication in host defense against infection we measured fly survival upon oral *P. aeruginosa* administration and CycE overexpression or downregulation in EBs. We found that CycE overexpression in EBs, which promotes their endoreplication to the expense of ISC mitosis (Fig. 3g-i), consistently improves fly survival (Supplementary Fig. 6a-b). On the other hand, CycE downregulation in EBs, does not inhibit endoreplication uniformly (Fig. 3j-l) and does not improve fly survival consistently (Supplementary Fig. 6a-b). We conclude that the cell autonomous induction of endoreplication in EBs by CycE promotes host defense to infection.

CycE downregulation in ISCs, which inhibits mitosis, also reduces fly survival, despite the increase in endoreplication (Supplementary Fig. 6c-d). This indicates that the trigger of endoreplication cannot fully compensate for the reduction of mitosis, and is in agreement with the higher survival rates of the highly mitotic strains that exhibited lower endoreplication (Suppl. Fig. 1).

Surprisingly, CycE overexpression in ISCs, which promoted mitosis to the expense of endoreplication in the anterior midgut (Fig. 3a-c), reduced fly survival (Supplementary Fig. 6c). Because *cycE* overexpression in ISCs using both the *Dl-Gal4* and the *ISC*^*ts*^*-Gal4* driver consistently reduced fly survival (Supplementary Fig. 6c-d), we suggest that the cell autonomous induction of mitosis in ISCs by CycE may lead to improper EC differentiation or function. This is supported by further data showing that *cycE* overexpression in ISCs increases dysplastic cell cluster formation, while *cycE* downregulation in ISCs decreases dysplasia (Fig. 4i). We conclude that both ISC mitosis and EB endoreplication may protect the *Drosophila* midgut from an infection and that improper EC differentiation may compromise fly resilience to intestinal pathogens.

### Highly mitotic strains are prone to dysplastic cell cluster formation

We next sought to assess whether highly mitotic strains, which cope better with a lethal intestinal infection (Suppl. Fig. 1), are prone to midgut dysplasia that normally occurs during aging or in genetically predisposed flies upon infection (Apidianakis, 2009; Biteau, 2008; Marianes, 2013, Siudeja 2015). We found that young flies of the highly mitotic strains develop more dysplastic cell clusters, that is, groups of 5 or more ISC-like cells or EEs upon infection and chemical inhibition of the Notch signaling pathway (Fig. 4a-c). Also, untreated flies of the highly mitotic strains develop more ISC-like and EE clusters at a young age (Fig. 4d-e) in agreement with their higher expression of the ISC marker *Delta* (*Dl*) (Guo, 2015) (Fig. 5a). Similarly, within an independent set of 4 additional genotypes, the mitosis level correlates with ISC-like/EE cluster incidence (Suppl. Fig. 7a-d). In addition, increasing or decreasing ISC mitosis directly through *cycE* overexpression or *cycE* downregulation in ISCs, respectively (Fig. 3a-b), alters the ISC-like/EE cluster incidence upon infection and Notch pathway inhibition according to the level of mitosis (Fig. 4i, j).

**Figure 5.**
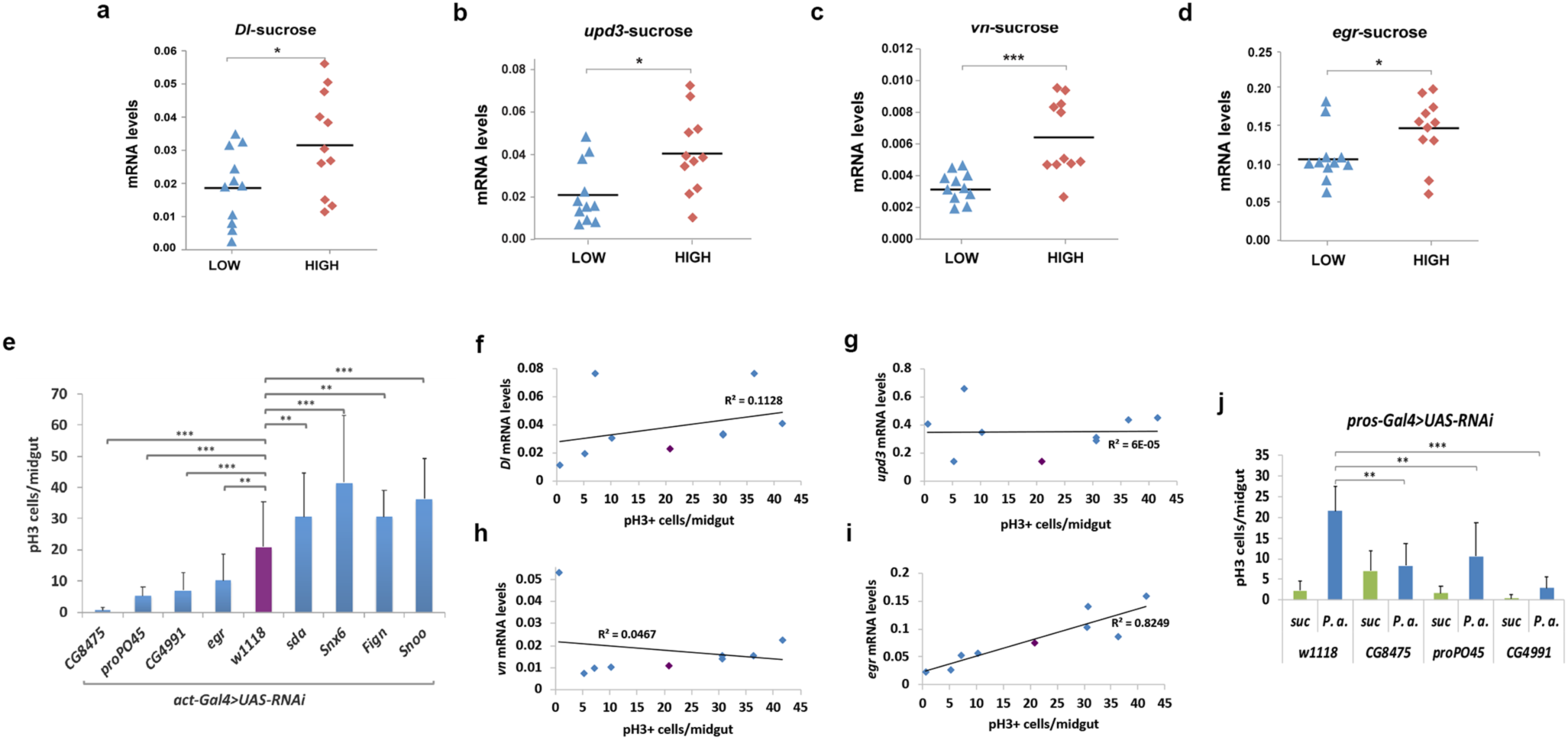
Differential expression of regenerative inflammation genes between the highly and the lowly mitotic strains. **a-d.** The 11 “High” vs. 11 “Low” mitosis strains differ in expression (t-test assessed) of *Dl* (a), *upd3* (b), *vn* (c) and *egr* (d). **e.** Mitotic cells per infected midgut for 8 polymorphism-associated genes identified via GWAS knocked-down ubiquitously (t-test assessed). **f-i.** Correlation between *Dl* (f), *upd3* (g), *vn* (h) and (i) *egr* expression and mitotic cells per midgut of the genotypes shown in e. **j.** Mitotic cells per midgut of flies in which *CG8475, proPO45* and *CG4991* are knocked down in the EEs using *pros*^*V1*^*-Gal4* (t-test assessed). ns *p*>0.05; * 0.01<*p*≤ 0.05; ** 0.001<*p*≤ 0.01; *** *p*≤ 0.001.

**Figure 6.**
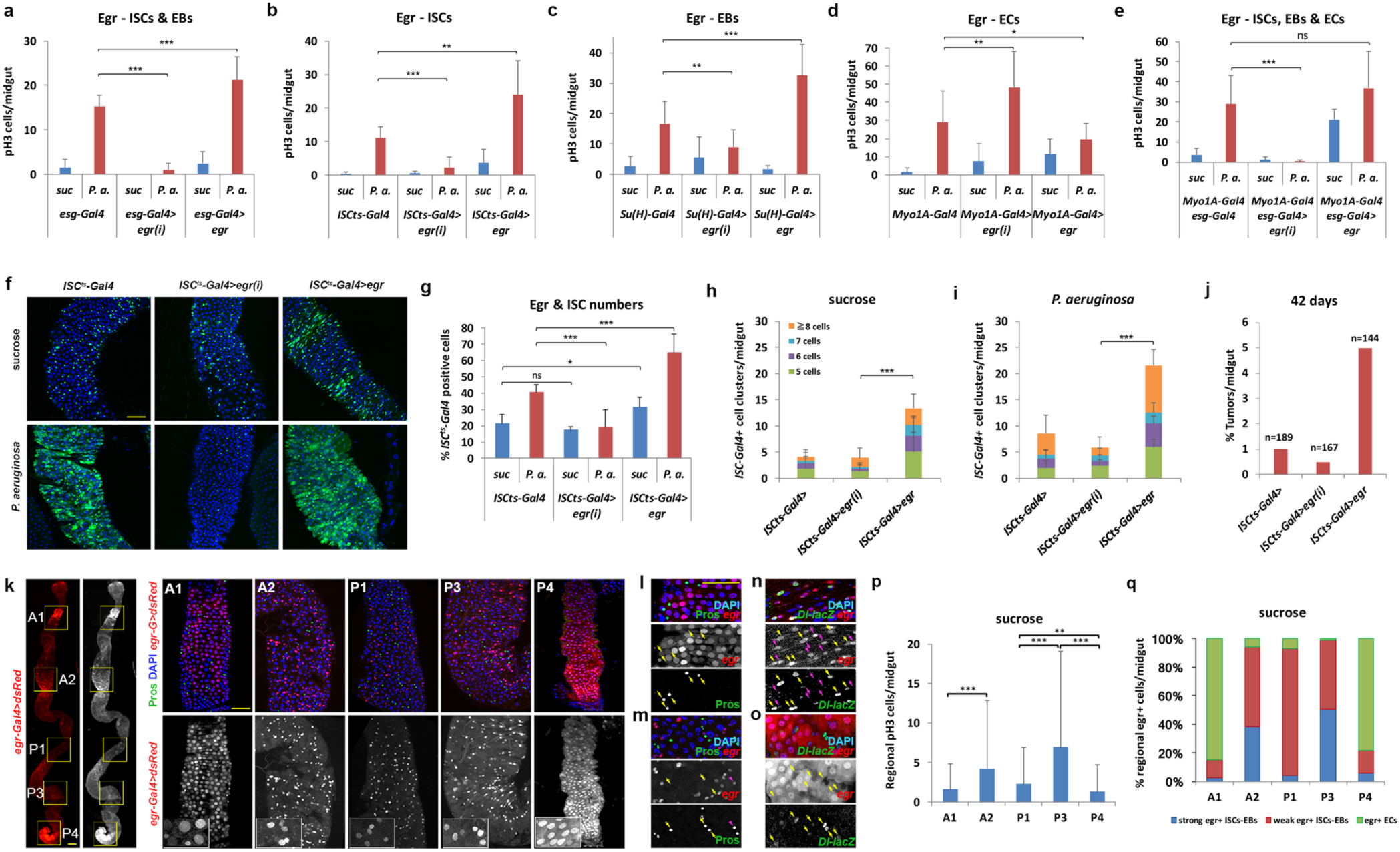
*egr* affects mitosis and dysplasia, and it is expressed in ISCs, EBs and ECs in different compartments of the *Drosophila* midgut. **a-e.** *egr* overexpression and *egr* RNAi in ISCs+EBs (a), ISCs (b), EBs (c), ECs (d), ISCs+EBs+ECs (e). **f-g.** *egr* RNAi and *egr* overexpression in ISCs decreases and increases ISC^ts^-positive cells in baseline and infected conditions as seen in images (f) stained with GFP (ISC^ts^) and DAPI (blue) and quantified (g). **h-j.** *egr* RNAi and *egr* overexpression in ISCs affects dysplastic ISC-like and EE cell cluster formation in baseline (h) and infected (i) conditions, as well as age-associated tumor incidence (j); h-i pairwise comparisons were chi-square tested. **k.** Distinct pattern of expression of *egr-Gal4 UAS-dsRed* stained with Prospero (green) along the midgut with prominent EC expression in A1 and P4 and strong progenitor expression in A2 and P3. **l-m.** Prospero (yellow arrows in single channel images) and *egr* (red) co-localize only rarely (purple arrowhead). **n-o.** Prominent but not exclusive co-localization of *Dl-lacZ* (green, yellow arrows in single channel images) with *egr* (red, purple arrows in single channel images). **p.** mitotic cells per A1, A2, P1, P3 and P4 region. **q.** Percentage of strongly and weakly expressing *egr* progenitors, and ECs expressing *egr* per A1, A2, P1, P3 and P4 region. Significance is assessed by t-test in panels a-e, g and p. ns *p*>0.05; * 0.01<*p*≤ 0.05; ** 0.001<*p*≤ 0.01; *** *p*≤ 0.001. Scale bars 50 um.

**Figure 7.**
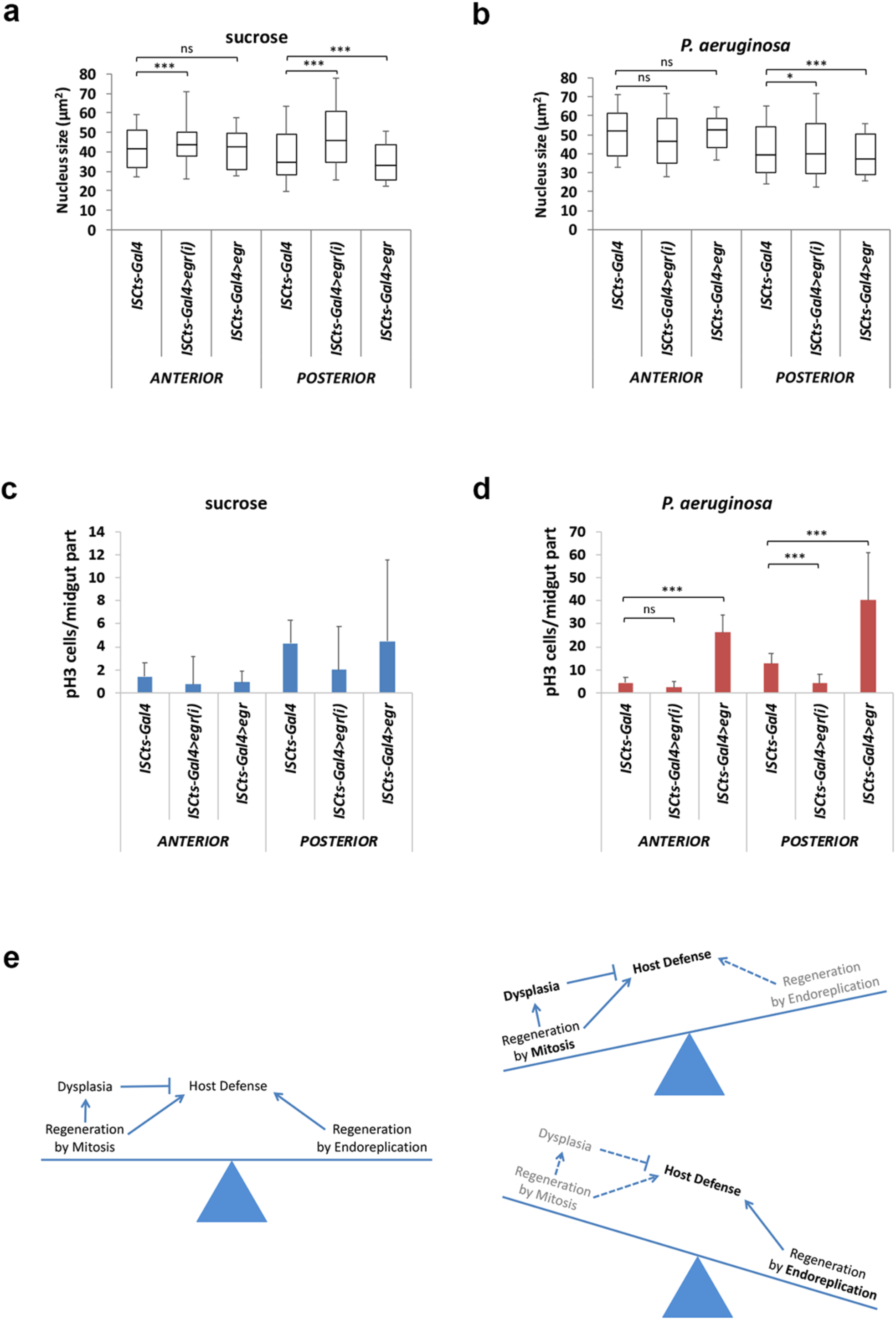
*egr* affects the mitosis-endoreplication balance in the midgut and signals via both Wgn and Grnd. **a-d.** *egr* RNAi and *egr* overexpression in ISCs non-autonomously affect EC endoreplication of the posterior midgut in baseline (a) and infected (b) conditions and the effects are inversely correlated with mitosis (c-d). **e.** Model of the intertwined trade-offs that control *Drosophila* midgut regeneration and host defense. ns *p*>0.05; * 0.01<*p*≤ 0.05; ** 0.001<*p*≤ 0.01; *** *p*≤ 0.001.

Moreover, 30- and 42-day old flies of the highly mitotic strains develop more ISC-like/EE clusters upon aging (Fig. 4f-h, 4m-o). Interestingly, the posterior midgut, which exhibits increased overall mitosis compared to the anterior, was more inflicted by ISC-like/EE dysplastic cell cluster (Fig. 4k-l, 4p-q). Thus, ISC mitosis is a key factor in promoting intestinal dysplasia in the form of dysplastic ISC-like/EE cell cluster formation that is widespread in young chemically-treated and infected flies, and progressive in flies upon aging.

### Egr and associated genes are linked to ISC mitosis

To explain mitotic variation at the molecular level, we undertook two approaches: (a) candidate gene expression assessment using real time quantitative PCR of the 22 strains exhibiting extreme mitosis upon infection, and (b) a genome wide association study (GWAS) using mitotic variation among all the 153 DGRP strains tested.

Assessing a list of 26 candidate genes related to *Drosophila* immunity or regeneration we identified the *Delta* (*Dl*), *unpaired 3* (*upd3*), *vein* (*vn*), and *eiger* (*egr*) genes encoding pathway ligands as highly expressed in the untreated highly mitotic strains (Table 1; Fig. 5a-d). These ligands are known stem cell signaling regulators, except from *egr* which is only recently linked to stem cell function (Guo, 2015; Jiang, 2009; Buchon, 2010; Doupe, 2018). Our GWAS analysis identified 95 variants, that is, single nucleotide polymorphisms (SNPs) and small insertions-deletions (INDELs) associated with midgut mitosis, corresponding to 39 protein-coding genes (Table 2). These genes could affect mitosis either locally in the migut or systemically. Ubiquitous downregulation for 8 of these genes was lethal to the flies, but downregulation of 7 of the rest of the genes plus *egr* reproducibly affected ISC mitosis. Downregulation with RNAi of CG8475, *proPO45*, CG4991 and *egr* reduced mitosis, while RNAi of *sda, Snx6, Fign* and *Snoo* increased mitosis (Fig. 5e). To identify potential cross-regulations between *Dl, upd3, vn* and *egr* and newly the above GWAS genes we correlated the expression of each of these ligands with the mitosis level of flies compromised in the expression of the 7 mitosis-related GWAS genes plus *egr* (Fig. 5f-i). We found that *egr* expression was the only one to positively and significantly correlate with the level of mitosis in these flies (Fig. 5i; *P*=00071). Moreover, based on FlyGut-seq data analysis *CG8475, proPO45* and *CG4991* are all enriched in EEs among all midgut cells (Buchon, 2013). Thus, we downregulated them specifically in the EEs via *prosV1-Gal4* and found that RNAi for all 3 genes consistently reduced mitosis in the midgut (Fig. 5j). Thus, *egr* and genes related to its expression have a key role of in ISC mitosis.

**Table 2:**
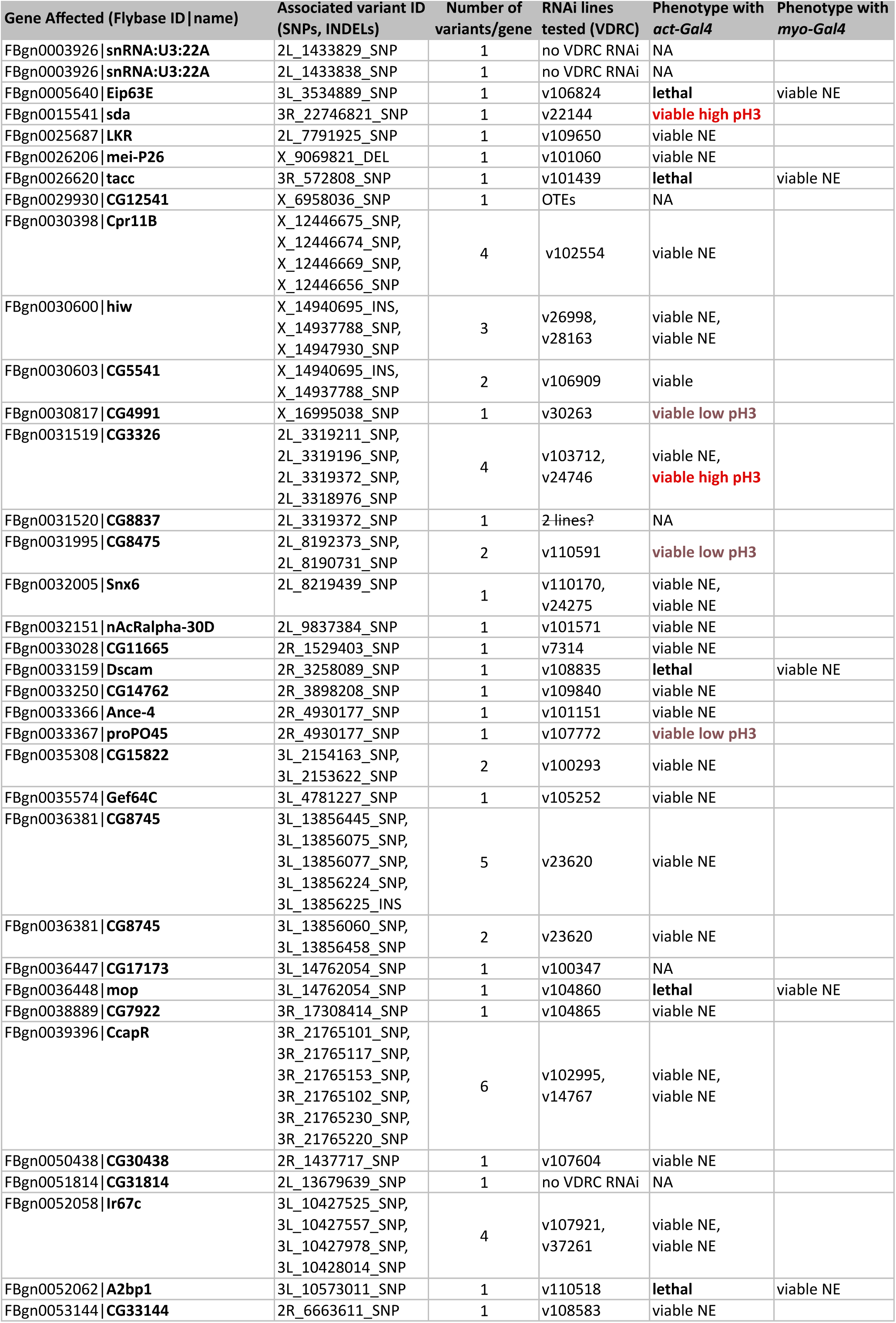
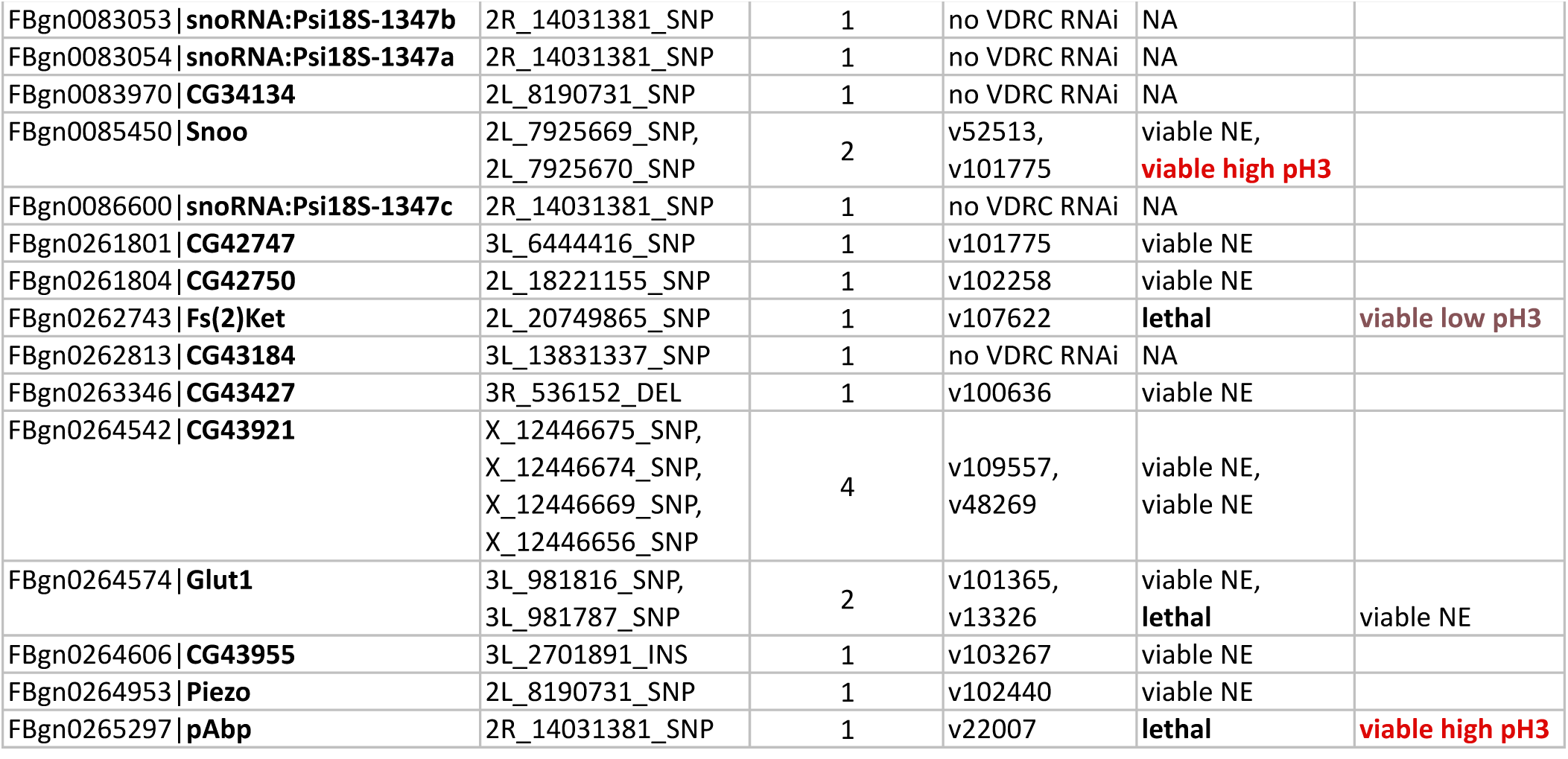
Genes identified by GWAS analysis. Genes associated with variants, the molecular nature of variants and their number for each gene are shown. Whole-body RNAi knockdown using the VDRC lines indicated for each gene was performed using the *act-Gal4* driver and intestinal mitosis was assessed. The 8 UAS-RNAi lines that did not produce viable offspring with *act-Gal4* were crossed to *Myo-Gal4*. NE: no effect.

### Tissue-intrinsic Eiger acts as an accelerator of mitosis and dysplasia to the expense of endoreplication

To identify the tissues and cell types in which *egr* is needed for ISC mitosis, we altered its expression systemically in the fat body and hemocytes, tissues previously shown to express *egr* and affect organismal metabolism and inflammation (Agrawal, 2016; Parisi, 2014; Mabery, 2010) (Suppl. Fig. 8). Surprisingly, while *egr* is induced in the fat body upon intestinal infection (Suppl. Fig. 8a-b), we did not observe any impact of systemic *egr* expression to ISC mitosis upon infection (Suppl. Fig. 8c-d).

To the contrary, we noticed a tissue-intrinsic role for *egr* by manipulating its expression levels either in all midgut epithelium cells (ISCs plus EBs plus ECs), or specifically in ISCs, EBs or ECs (Fig. 6a-e). We find that downregulation of *egr* in all midgut epithelium cells eliminates infection-induced ISC mitosis, whereas *egr* overexpression is sufficient to increase mitosis even in the absence of infection (Fig. 6e). Moreover, *egr* overexpression in ISCs, EBs or ISCs+EBs significantly induced ISC mitosis upon infection, and *vice versa egr* downregulation in ISCs, Ebs or ISCs+EBs significantly reduced ISC mitosis upon infection (Fig. 6a-c), while *egr* overexpression only in ISCs without infection only marginally induced mitosis, indicating that *egr* secreted from ECs may contribute to ISC mitosis (Fig. 6b). Accordingly, *egr* overexpresion in the ECs induced ISC proliferation at baseline, but surprisingly it inhibited proliferation upon infection (Fig. 6d). In terms of stem cell numbers, overexpression of *egr* in ISCs using *ISC*^*ts*^*-Gal4 UAS-GFP* increased the GFP positive cells at baseline and upon infection (Fig. 6f-g). Moreover, *egr* overexpression via *ISC*^*ts*^*-Gal4 UAS-GFP* induces dysplastic cell clusters both in the presence and the absence of infection (Fig. 6h-i) and promoted spontaneous tumor formation upon aging (Fig. 6j). Thus, intrinsic expression of *egr* in ISCs and other epithelial cells increases their proliferation and predisposition for tumorigenesis.

Using *egr-Gal4 UAS-dsRed* flies, we noticed *egr* expression in the ISCs and EBs along the whole midgut, but also in the ECs in two low mitosis zones (the A1 and P4) (Fig. 6k-o). *egr* rarely co-localizes with Prospero expression (1.5% of *egr*+ cells upon infection, n=530; and 1.8% without infection, n=550), which labels the EEs and their precursors (Zeng, 2015; Guo, 2015). In addition, we used an Egr-GFP protein trap line (Sarov, 2016) and found that Egr is localized in the cytoplasm of intestinal progenitors and ECs, but not EEs (Supp. Fig. 9). Interestingly, we noticed that mitosis spikes in the A2 and the P3 regions of the midgut, the exact same regions with the highest progenitor (ISCs+EBs) to enterocyte (EC) ratio of *egr* expression (Fig. 6p-q). Of note, *egr* expression was not induced upon infection in the midgut (Suppl. Fig. 10a). And, Ras signaling induction in ISCs, which is necessary and sufficient for regeneration upon infection (Jiang, 2011), does not induce *egr* (Suppl. Fig. 10b). We conclude that *egr* expression is inherently controlled in the midgut progenitors through baseline signals rather than those induced by upon infection.

To assess the role of *egr* in the interplay between midgut mitosis and endoreplication we downregulated and overexpressed *egr* in ISCs at baseline and upon infection. *erg* RNAi led to an increase in EC endoreplication at baseline and upon infection in the anterior and posterior midgut (Fig. 7a-b). To the contrary, *erg* overexpression led to a decrease in EC endoreplication at baseline and upon infection in the posterior midgut (Fig. 7a-b), while mitosis was increased upon infection in the same midgut region (Fig. 7c-d). Thus, *egr* expression in ISCs may directly promote mitosis in ISCs and dysplasia to the expense of EC endoreplication at a given homeostatic demand.

## Discussion

Similarly to mammals, *Drosophila* has evolved two mechanisms to maintain tissue integrity upon injury: compensatory cell proliferation and compensatory cellular overgrowth of differentiated cells (Tamori, 2014; Huh, 2004). While the *Drosophila* midgut responds to injury by deploying both mechanisms at the same time, little is known on how new cell supply by ISC mitosis and EC growth by endoreplication are coordinated during homeostasis. Homeostasis can be achieved not only at baseline in the absence of extensive tissue damage, but also during infection where the cell loss is balanced by new or bigger cells. Our studies on the *Drosophila* midgut provide evidence that the differentiating ECs undergo more cycles of endoreplication as a means to increase their size and sustain tissue integrity and dimensions, when mitosis is limited. We show that the level of mitosis per fly midgut can differ significantly from strain to strain at the baseline or during intestinal infection; and strains exhibiting high stem cell mitosis have increased midgut length and posterior width. Nevertheless, during infection both high and low mitosis strains increase their anterior and posterior width, but not their midgut length. The increase in midgut width upon infection is accompanied by an increase in endoreplication in the anterior and posterior midgut in the low rather than the high mitosis strains. While preserving adult midgut width primarily via endoreplication, lowly mitotic strains cope with bacterial infection less successfully than highly mitotic ones. Thus, endoreplication may provide a mechanism of tissue recovery from EC damage that is less affective than mitosis-driven cell renewal.

To corroborate the role of endoreplication in host defense to infection we altered the expression levels of *cycE* in either ISCs or EBs, and noticed that CycE induces ISCs to proliferate and EBs to endoreplicate cell-autonomously, but counterbalances these alternative cell cycle programs non-cell-autonomously. Importantly though boosting endoreplication in the EBs via cycE overexpression consistently improves survival to infection. On the other hand, boosting mitosis in the ISCs via *cycE* overexpression consistently reduces host survival to infection due to the ensuing dysplasia i.e. the abundance to ISC-like cells and EE cell clusters. Host survival is also reduced upon *cycE* RNAi in the ISCs. Therefore, both ISC mitosis and EB endoreplication are necessary for optimal host defense to infection, and that excessive mitosis causes dysplasia and compromised host defense. In this respect, endoreplication may facilitate a primary response to injury or increased nutrient availability, when both mitosis and endoreplication need to be deployed to restore tissue integrity in a timely manner. At a given cell demand, a balance between cell loss and cell replacement or growth can be achieved either with high mitosis and low endoreplication or vice versa with low mitosis and high endoreplication. This is terms with previous studies showing that injury and nutrient increase deploy endoreplication as a faster or supporting means to achieve homeostasis (Buchon, 2010; Choi, 2011; Losick, 2013).

On the other hand, the level of mitosis, as opposed to endoreplication, correlates positively with tissue dysplasia. Highly mitotic midguts develop clusters of mis-differentiated ISCs expressing the ISC marker *Delta* and EE clusters expressing Prospero, upon damage and inhibition of *Notch*, but also spontaneously during aging. Although less prevalent, ISC-like and EE clusters are prominently formed in untreated highly mitotic strains in agreement with their higher expression of the ISC marker *Delta*. Accordingly, increasing ISC mitosis directly through *cycE* overexpression in ISCs increases ISC-like and EE cluster incidence. And *vice versa* decreasing ISC mitosis directly through *cycE* downregulation in ISCs decreases ISC-like and EE cluster incidence. Thus, high mitosis may boost ISCs to produce more dysplastic cells once these are formed or impedes normal differentiation of ECs causing dyspastic cell formation.

Our quantitative genetics study demonstrates that *Drosophila* strains exhibiting excessive mitosis upon infection are characterized by higher baseline mitotic activity and increased baseline expression of regenerative inflammation genes. We pinpoint four key signaling pathway ligands in controlling high level of mitosis: the homeostasis ISC marker Delta (Dl), the stress-induced cytokine Upd3 emanating from ECs, the visceral muscle-emanating growth factor Vein and the cytokine Egr, which is expressed in most ISCs and EBs (Duppe 2018). Mild *egr* expression in ISCs may not suffice to induce mitosis in the absence of a secondary stimulus, but stronger *egr* expression and or expression from adjacent cell does. Evidently though, *egr* expression in ISCs consistently and significantly facilitates ISC mitosis upon infection. Thus, Egr is another inflammatory regeneration cytokine along with Upds that acts not only systemically or as part of the host immunity, but also to regenerate the midgut epithelium. Regenerative inflammation can be induced upon infection and cell stress via STAT signaling in flies and mice (Jiang, 2009; Taniguchi, 2012), and our work adds to this knowledge providing another example of a mitosis promoting cytokine that is constantly and tissue-intrinsically expressed in the intestine. Midgut *egr* expression is not induced by infection or upon Ras signaling in ISCs, which is key for inducing mitosis (Jiang, 2011). Nevertheless, our GWAS analysis indicates that the downregulation of CG8475, *proPO45*, CG4991 correlates with a lower *egr* expression and a lower mitosis as a response to infection. Downregulation of these three genes specifically in EEs, where they are more abundant among other midgut cells (Dutta 2015), suffices to reduce mitosis. Thus, it is intriguing to study such signals that sustain *egr* expression in the midgut at baseline making ISCs more responsive to infection.

Similarly to the intestinal epithelium of *Drosophila* the mammalian one is maintained by ISCs that divide to give rise to ISCs and transient progenitors, which normally differentiate into ECs, EEs and other cells (Zeng, 2015; Guo, 2015). Given the impact of ISC microenvironment in controlling ISC mitosis (Maeda, 2008; Choi, 2011; Patel, 2015), we suggest that a regenerative inflammation signaling through Egr and other mitogens and cell growth factors may be fine-tuned pharmacologically or through diet and microbiota as a means to optimize regenerative capacity and tissue resilience to infection, while decreasing predisposition for dysplasia.

---

**Supplementary Figure 1.**
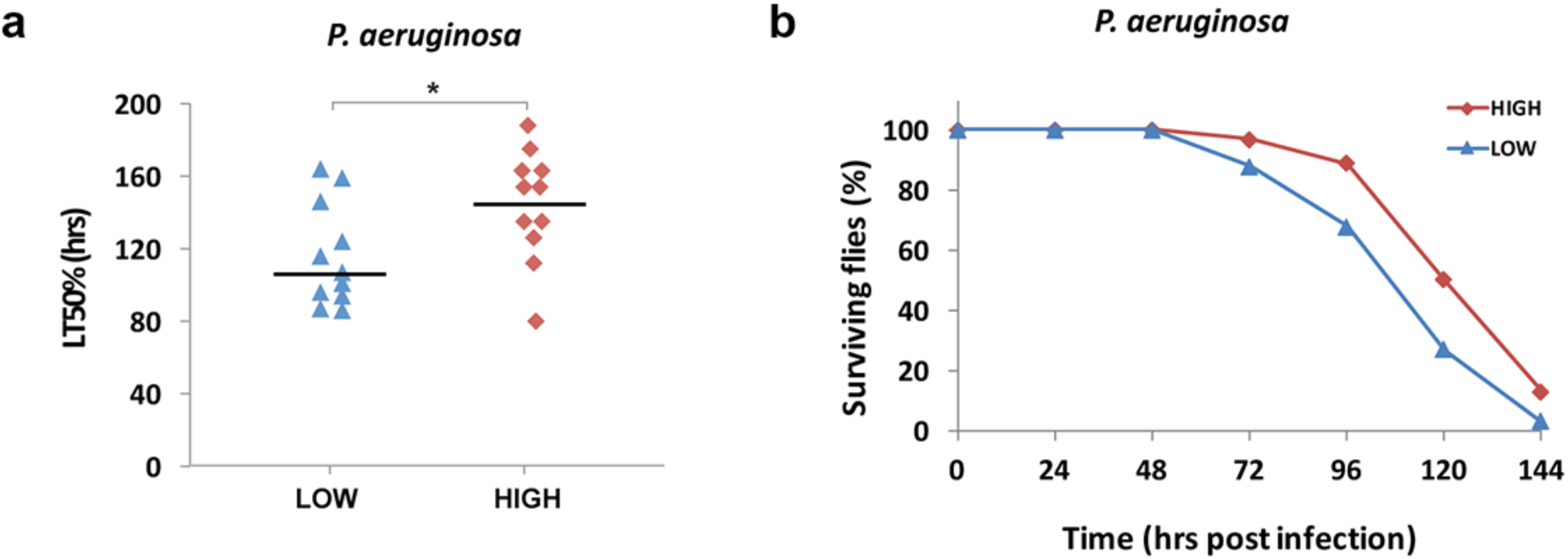
High mitosis strains survive longer upon *P. aeruginosa* infection. **a.** Individual assessment of the 22 extreme strains followed by plotting and statistical analysis of the lethal time 50% (LT50%) of the group of high vs. the group of low mitosis strains. t-tested * 0.01<*p*≤ 0.05. **b.** Pooled strain survival with 5 flies from each of the 22 strains forming a cohort. High mitosis strain flies are marked with *Dl-Gal4 UAS-scrGFP* (isogenized) to be distinguished from low mitotic ones and monitored for the length of their life. Statistical significance in b was *p*≤ 0.001.

**Supplementary Figure 2.**
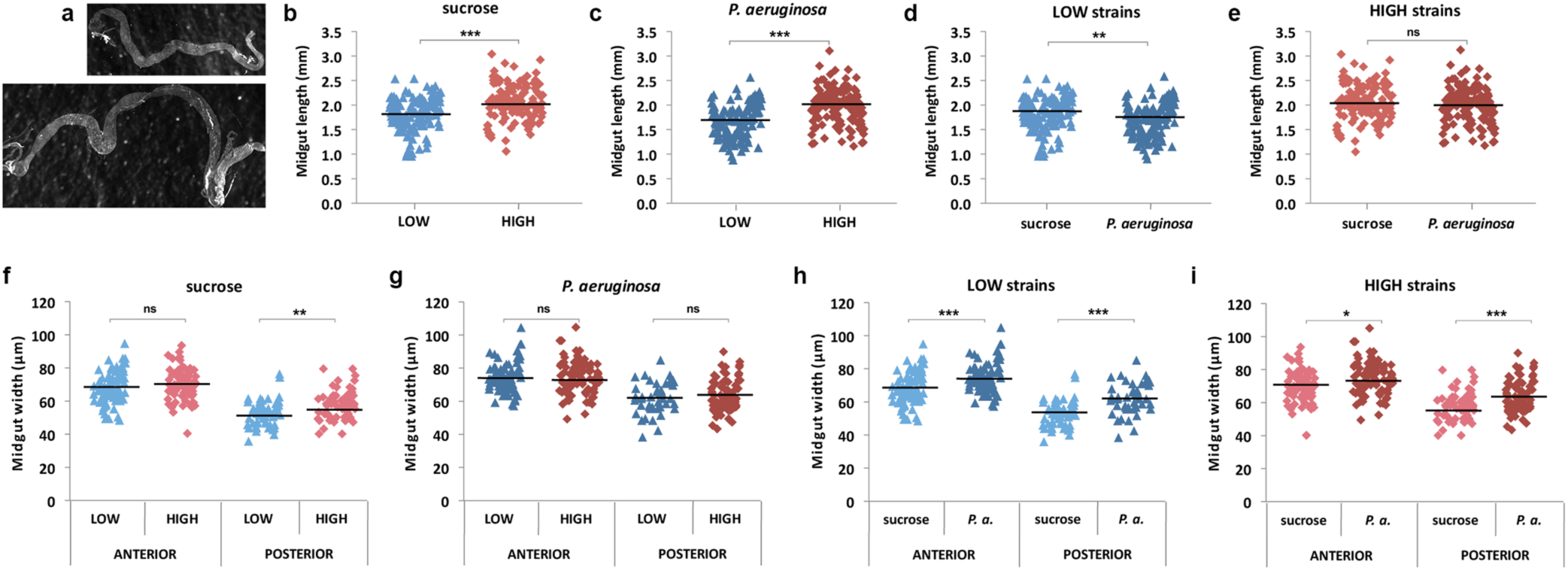
High mitosis strain midguts are larger in size, but do not respond to infection differently from low mitosis strains. **a.** A “low” and a “high” strain differing in length. **b-c.** Midgut is on average longer in “high” strains both before (b) and upon (c) infection. **d-e.** “Low” and “high” strains tend to lose length upon infection. **f-g.** Posterior midguts of the “high” strains are significantly wider than “low” before infection (f) and tentatively so upon infection (g). **h-i.** Both “low” (h) and “high” (i) strains increase the width of their anterior and posterior midgut upon infection. Statistical significance is assessed by t-test in b-i: ns *p*>0.05; * 0.01<*p*≤ 0.05; ** 0.001<*p*≤ 0.01; *** *p*≤ 0.001.

**Supplementary Figure 3.**
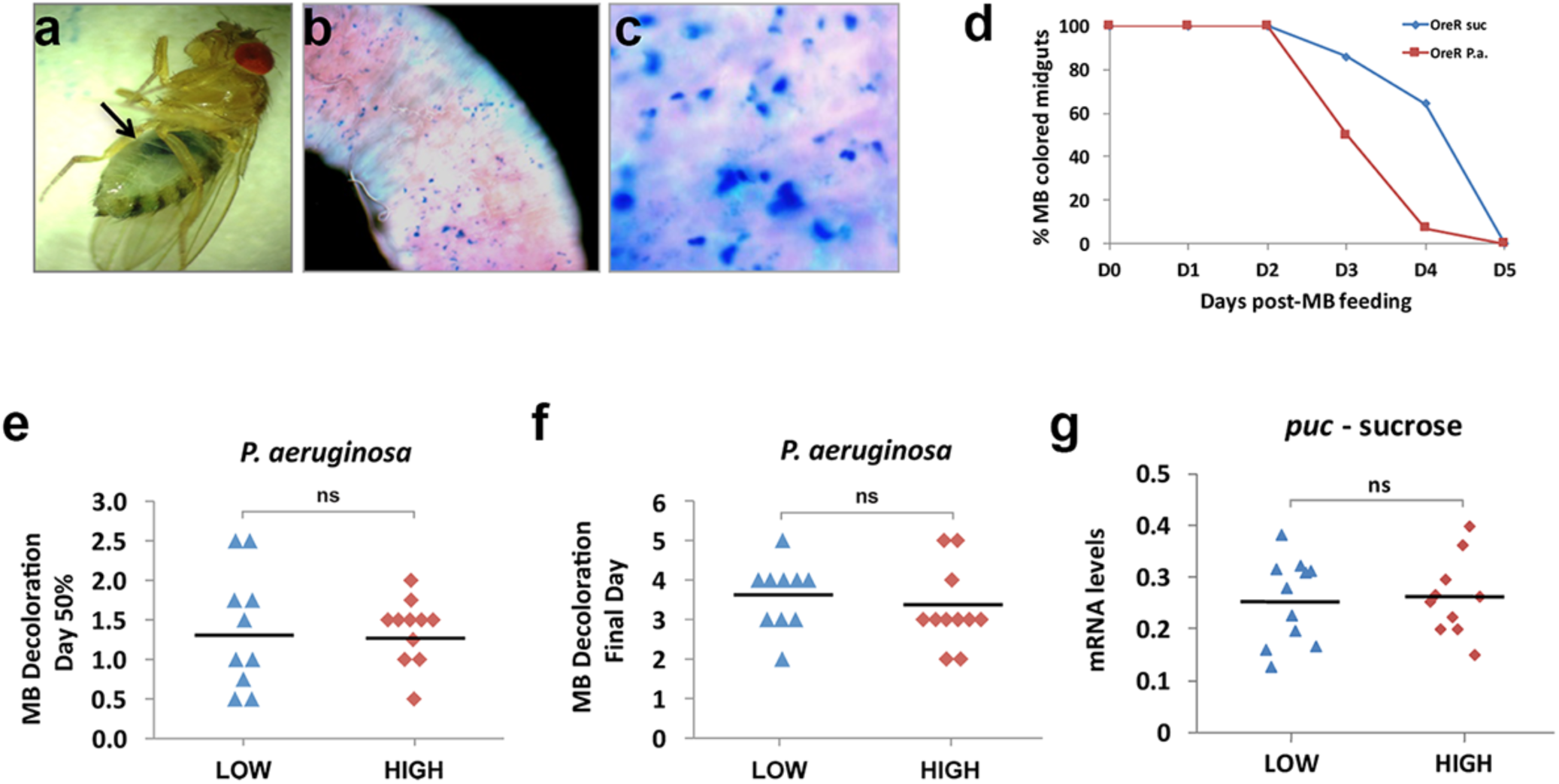
Midgut epithelium exfoliation and stress signaling response to infection are comparable between high and low mitosis strains. **a-c.** Methylene blue colored intestines as seen from the outside of the fly abdomen (a), in a part of a dissected midgut (b) and in higher magnification (c). **d**. Decoloration (loss of methylene blue coloration) is accelerated by infection. **e-f.** The day 50% (e) or 100% (f) of stained flies lose their color is similar between “high” and “low” strains. **g**. JNK pathway target *puckered* (*puc*) is similarly expressed between “high” and “low” strains. No statistical differences are observed in e-g (t-tested).

**Supplementary Figure 4.**
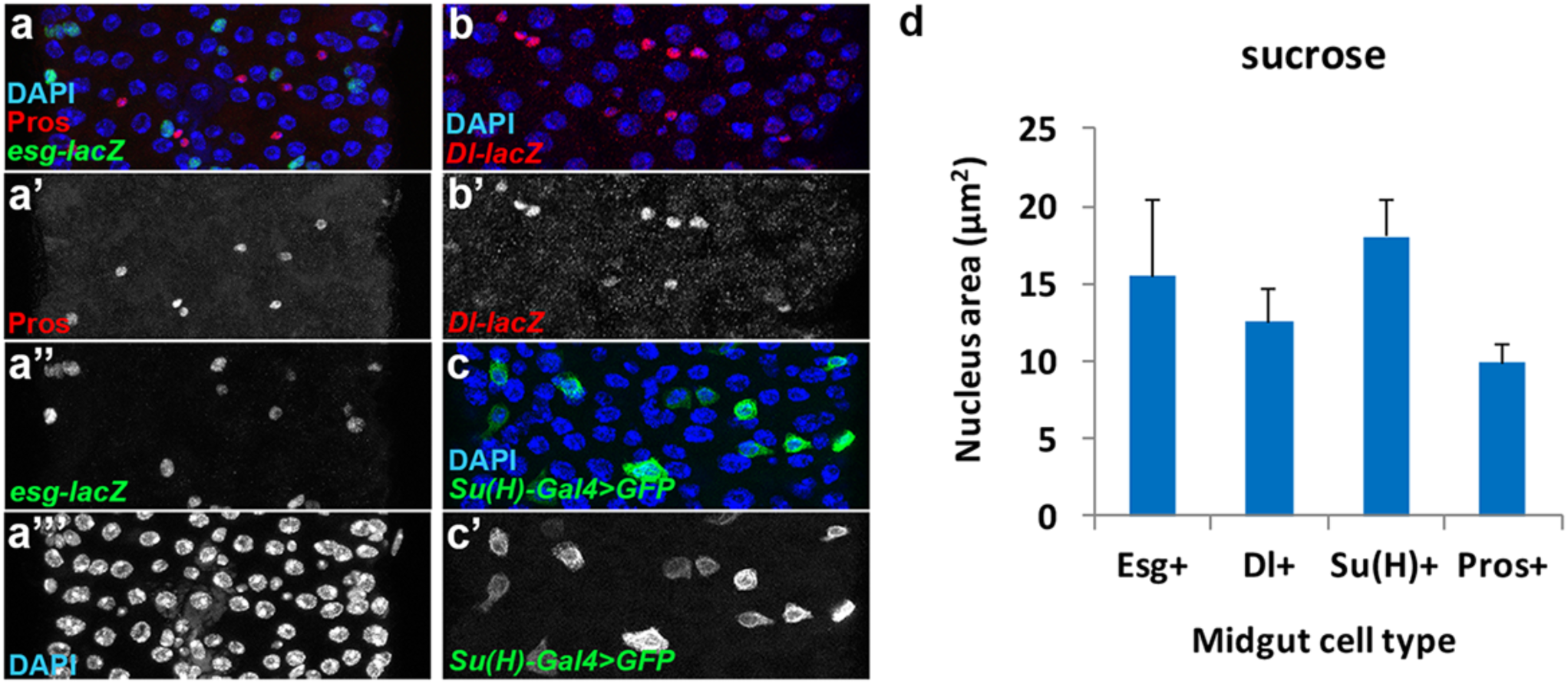
Nucleus size determination of the different midgut cell types. **a.** Midguts of the *esg-lacZ* enhancer trap were stained with anti-beta-Gal (green) and anti-Prospero (red) to label intestinal progenitors (ISCs and EBs) and EEs, respectively. **b.** Midguts of the *Dl-lacZ* enhancer trap were stained with anti-beta-Gal (red) to specifically label the ISCs. **c.** The *Su(H)-Gal4 UAS-GFP* line was used to specifically label the EBs (green). DAPI (blue) stains all nuclei in a-c. **d.** Quantification of nucleus surface of different midgut cell types from fluorescent images as in a-c.

**Supplementary Figure 5.**
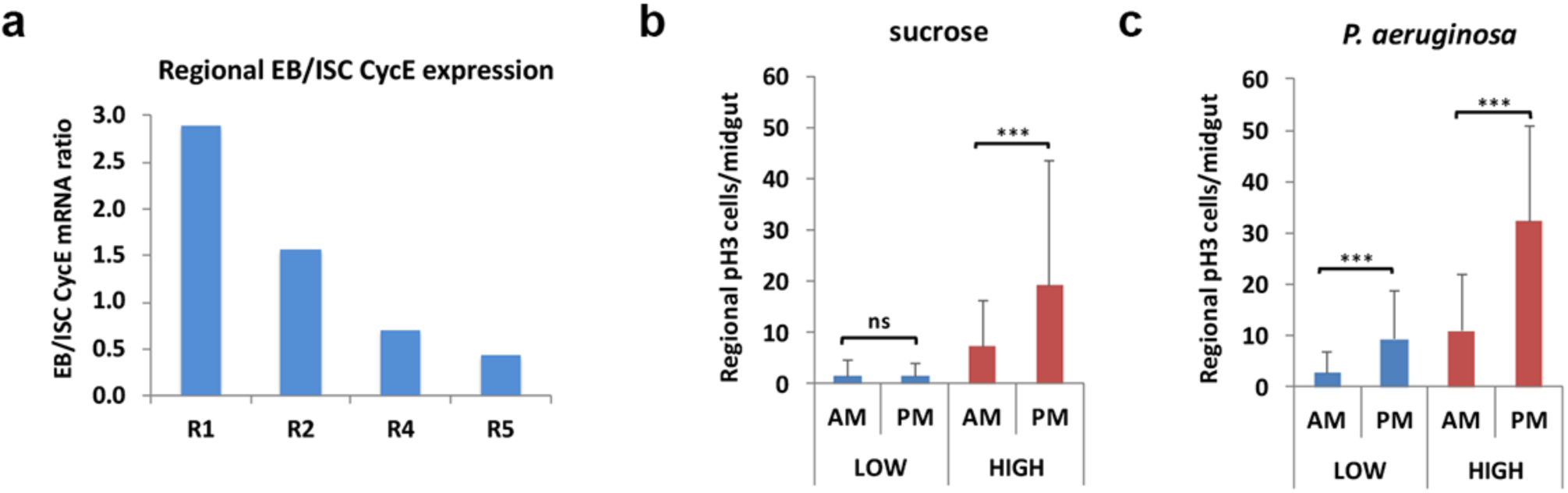
*cycE* expression is enriched in EBs compared to ISCs of the anterior midgut and this correlates with less mitosis. **a.** The ratio of EB/ISC *cycE* mRNA levels is plotted for the anterior (R1 and R2) and the posterior (R4 and R5) regions of the midgut (data from Buchon et al, 2013). **b-c.** Regional mitosis in low and high mitosis strains in baseline (b) and infected (c) conditions. Statistical significance is assessed by t-test in b-c: ns *p*>0.05; * 0.01<*p*≤ 0.05; ** 0.001<*p*≤ 0.01; *** *p*≤ 0.001.

**Supplementary Figure 6.**
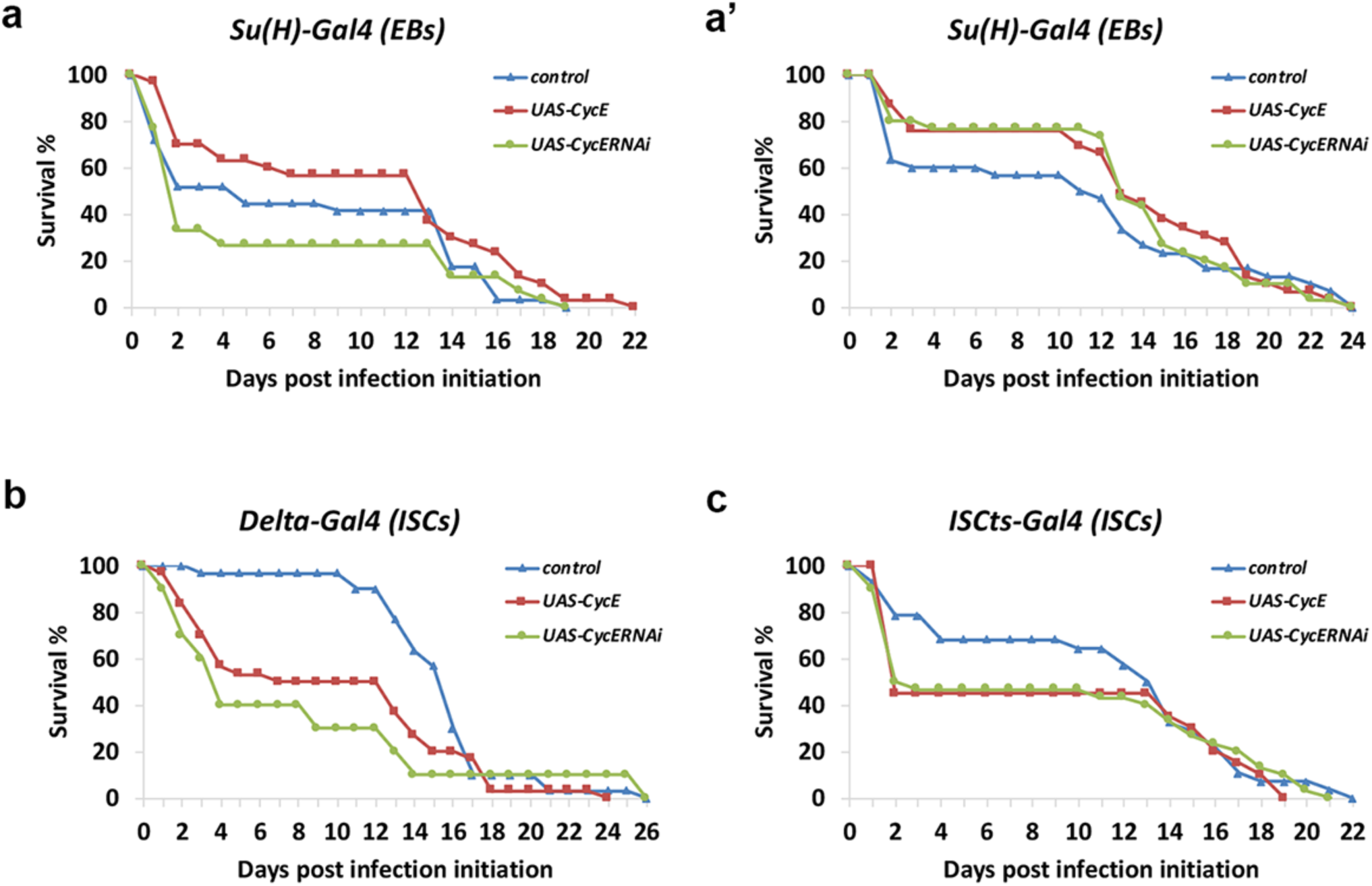
Manipulation of *cycE* levels in EBs and ISCs affects survival. **a-a’.** Survival of flies overexpressing *cycE* or *cycE*^*RNAi*^ in EBs upon *P. aeruginosa* infection. Two independent experiments are shown. Contrary to inconsistent effects of *cycE*^*RNAi*^, note consistent increase of survival upon *cycE* overexpression. **b-c.** Survival of flies overexpressing *cycE* or *cycE*^*RNAi*^ in ISCs upon *P. aeruginosa* infection using the drivers *ISC*^*ts*^*-Gal4* (b) and *Dl-Gal4* (c). Statistical significance in b and c was *p*≤ 0.001.

**Supplementary Figure 7.**
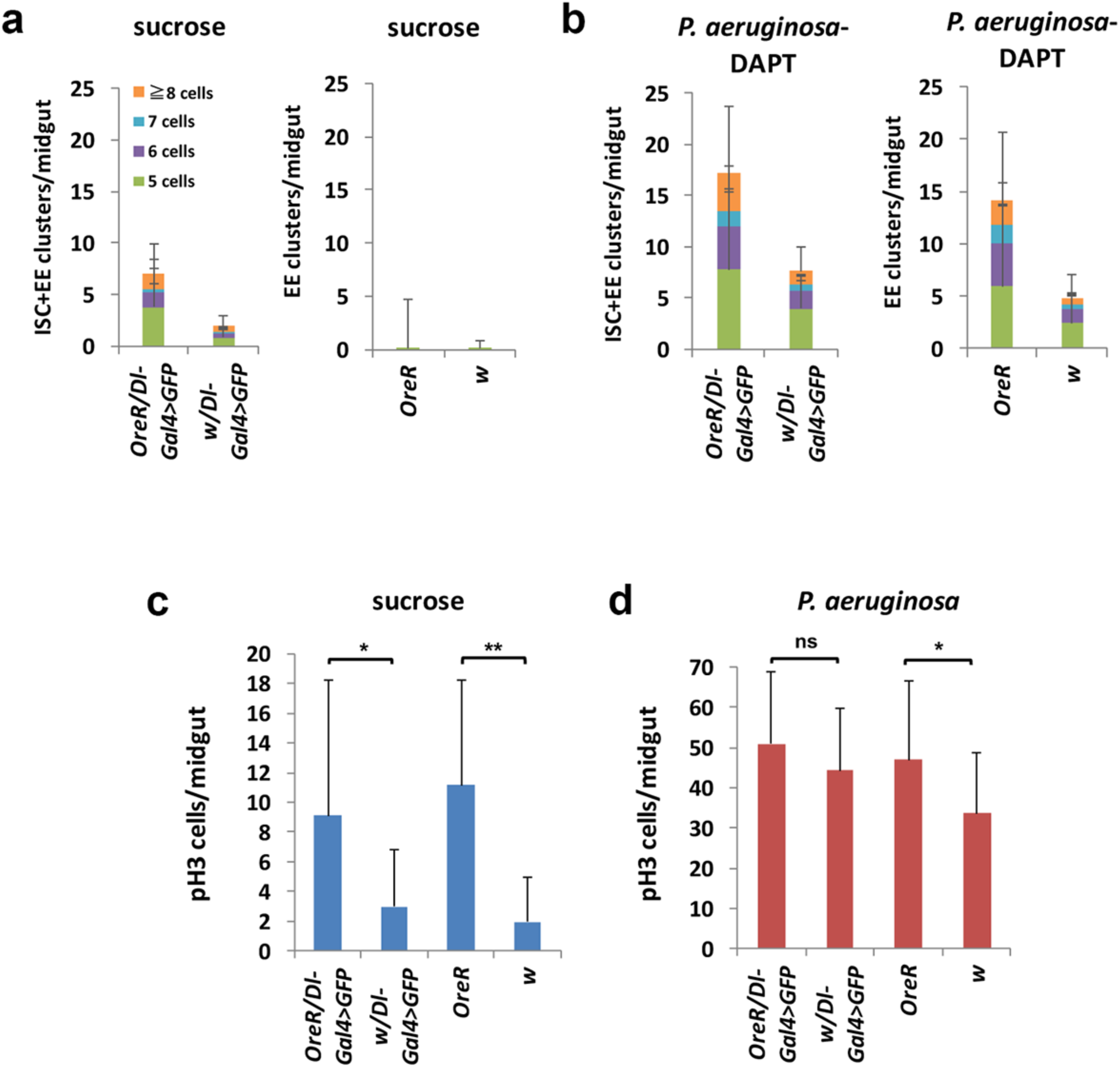
ISC-like and EE dysplastic cell clusters correlate with mitosis levels in different genotypes. **a-b.** Prospero and *Dl-Gal4 UAS-srcGFP* cell clusters of 5, 6, 7 and ≥8 cells without (a,) and upon DAPT/infection (b) Oregon R x *Dl-Gal4 UAS-srcGFP* and *w*^*1118*^ x *Dl-Gal4 UAS-srcGFP*, Oregon R (OreR) and *w*^*1118*^ (*w*). **c-d.** Midgut mitosis without (c) and with (d) infection in the genotypes assessed in a-b. Statistical significance is assessed by t-test in c-d: ns *p*>0.05; * 0.01<*p*≤ 0.05; ** 0.001<*p*≤ 0.01; *** *p*≤ 0.001.

**Supplementary Figure 8.**
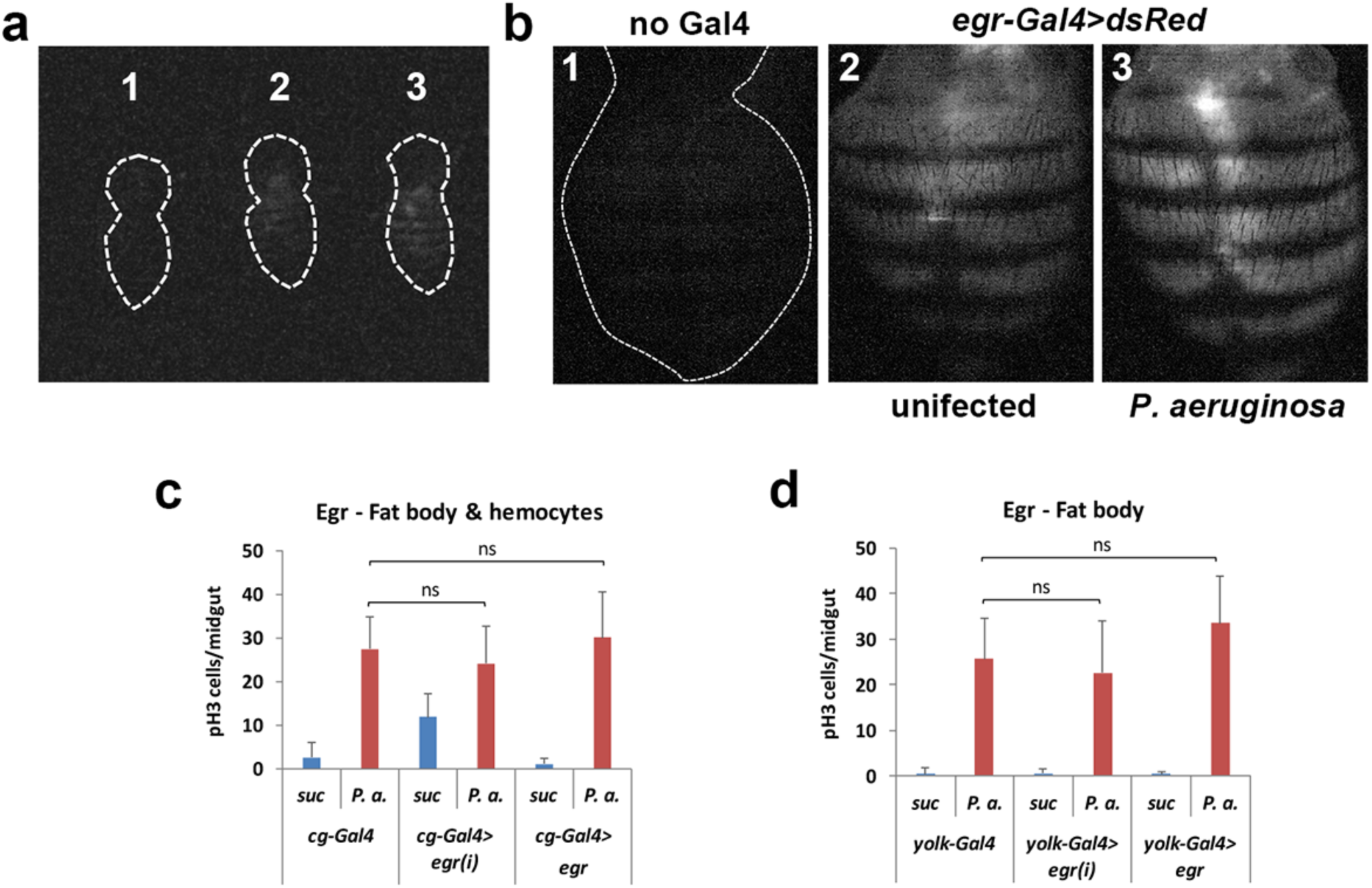
*egr-Gal4 UAS-dsRed* is expressed in the fly abdomen and while inducible upon infection, it does not contribute to midgut mitosis upon oral infection. **a-b.** Fluorescence of whole fly (a) and abdominal only (b) magnification of dorsal abdomen and imaging under the same settings of a fly without *egr-Gal4 UAS-dsRed* (1), an uninfected *egr-Gal4 UAS-dsRed* fly (2) and an orally infected *egr-Gal4 UAS-dsRed* fly (3). **c-d.** *egr* overexpression and *egr* RNAi in the fat body and hemocytes (c) or the fat body exclusively (d) of female flies do not contribute to midgut mitosis upon oral infection. No statistical differences are observed in c-d (t-tested).

**Supplementary Figure 9.**
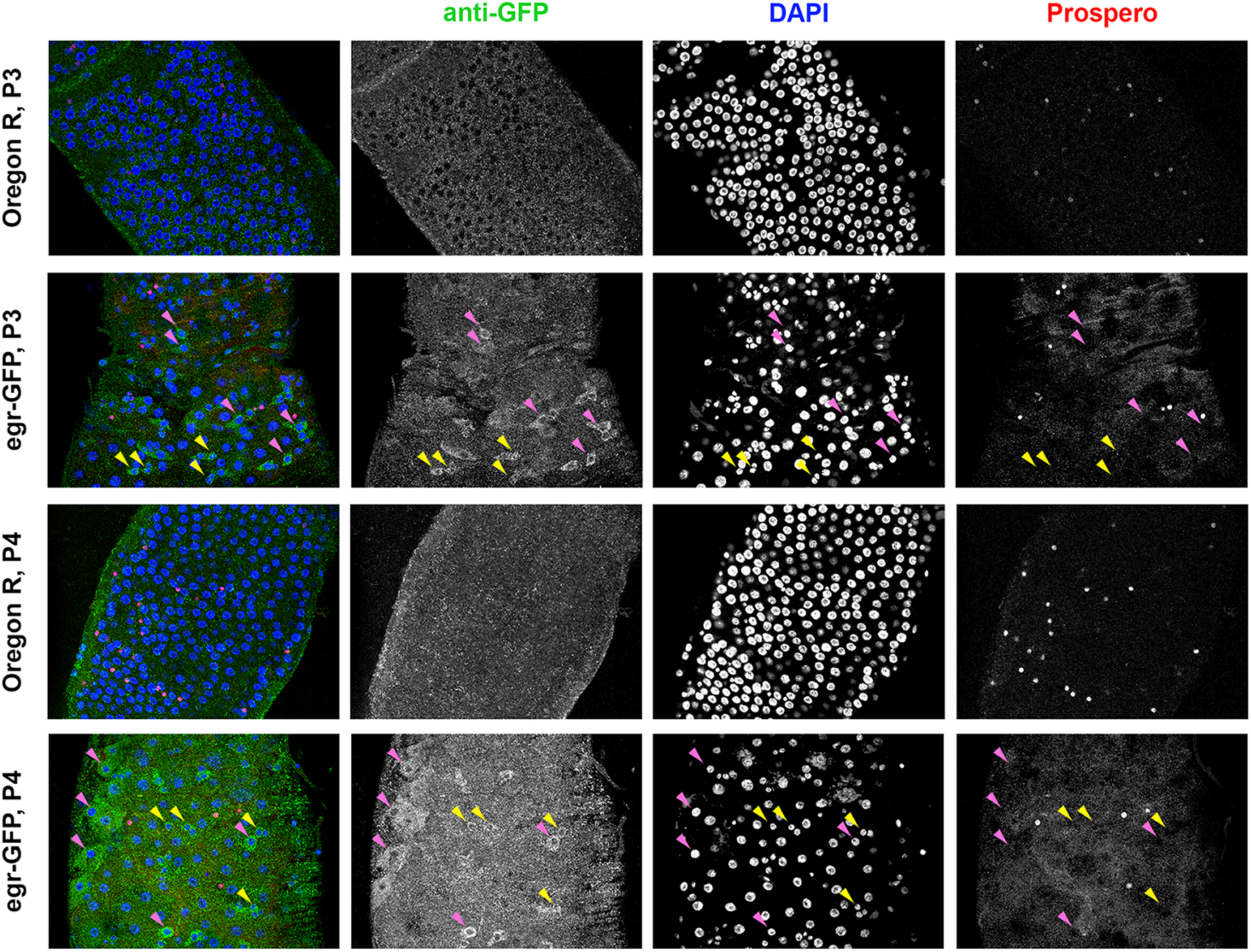
An Egr-GFP protein trap is expressed in intestinal progenitors and ECs of the adult posterior midgut. Confocal images of posterior midguts (regions P3 and P4) of Oregon R (control) and Egr-GFP flies immunostained with an antibody against GFP (green) and anti-Prospero (red). Egr-GFP labels progenitor cells (yellow arrowheads; cells with small nuclei negative for Prospero) and ECs (purple arrowheads; cells with polyploid nuclei).

**Supplementary Figure 10.**
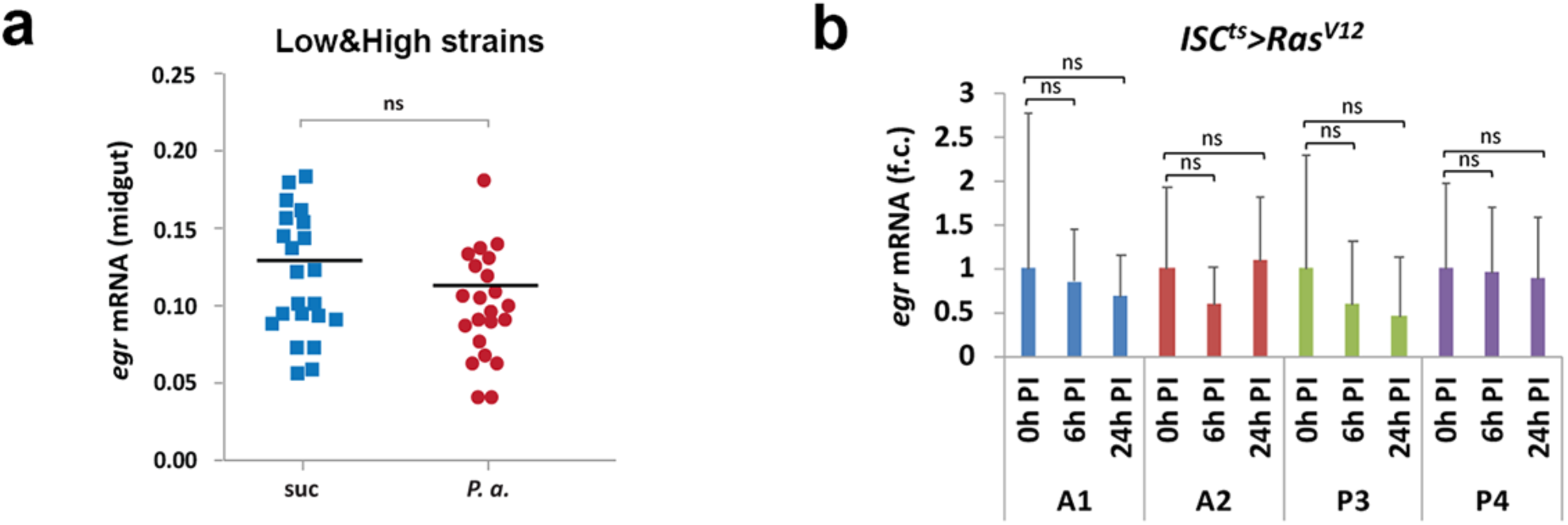
Infection and Ras signaling do not induce *egr* expression. **a-b.** *Pseudomonas aeruginosa* infection of high and low mitotic strains (a), and induction of Ras signaling in ISCs (b) do not induce *egr* expression in the midgut (regions A1, A2, P3 and P4 were tested independently in b). No statistical differences are observed in a-b (t-tested).

## Material and Methods

### Fly stocks

All stocks were maintained at 25°C on a 12:12 hour light:dark cycle on fly food containing yeast, cornmeal, sugar and agar supplemented with Tegasept and Propionic acid. The 153 inbred strains of the *Drosophila* Genetic Reference Panel (DGRP) were obtained from the Bloomington *Drosophila* Stock Center (BDSC). The 48 *UAS-RNAi* lines of genes selected through GWAS analysis (see Table 2) were obtained from the Vienna *Drosophila* RNAi stock Center (VDRC). 40 of them produced viable progeny when crossed to *w; actin-Gal4 UAS-GFP/CyO*. The rest were crossed to *w; Myo1A-Gal4 UAS-EGFP/CyO*. All UAS-RNAi lines were isogenized by backcrossing to *w*^*1118*^ for 6 generations.

The following Gal4 lines were used for tissue and cell-type specific expression: For EC expression: *tub-Gal80*^*ts*^*/FM7; Myo1A-Gal4 UAS-EGFP/CyO* (ref.11; Apidianakis et al PNAS 2009). For progenitor expression: *w; esg-Gal4 UAS-GFP tub-Gal80*^*ts*^ (ref.11; Apidianakis et al PNAS 2009). For EB expression: *w; Su(H)-Gal4 UAS-CD8GFP tub-Gal80*^*ts20*^*/CyO* (ref.28 Zeng et al Genesis 2010). For ISC expression: *w; UAS-src-GFP/CyO; Delta-Gal4 tub-Gal80*^*ts*^*/TM6C* (ref.29; Zeng et al Genesis 2010) and *esg-Gal4 UAS-GFP; Su(H)-Gal80 tub-Gal80*^*ts*^ (ref.19; Zeng & Hou Development 2015). For VM expression: *w; how24B-Gal4 UAS-EGFP/TM3* (originating from BDSC#1767) and *w; dmef2-Gal4 UAS-dsRed/TM3* (ref. 30; G. Ranganavakulu, Dev. Biol., 1996) were combined with *tub-Gal80*^*ts*^ on the X. For EC and progenitor expression: *w; esg-Gal4 UAS-GFP Myo1A-Gal4* (recombined on the 2^nd^ chromosome). For fat body expression: *w; yolk-Gal4* (gift from Norbert Perrimon). For fat body and hemocyte expression: *w; cg-Gal4* (BDSC#7011). *w; UAS-egr*^*RNAi*^ (VDRC# 108814 KK), *w; UAS-CycE*^*RNAi*^ (VDRC# 110204 KK), *w; UAS-CycE* (BDSC#4781), *UAS-Notch*^*IC5*^ (ref. 31; Go, et al Development 1998), *UAS-p53* (BDSC #8420). The following stocks were obtained from Pierre Léopold (ref.20; Andersen et al., 2015): *w; UAS-egr*^*strong*^, *w; UAS-wgn*^*IR*^, *w; UAS-grnd*^*extra*^*/TM6B, w; UAS-grnd*^*intra*^*/TM6B*. *w*^*1118*^ was used as control to UAS strains and *Oregon R* as a typical wild-type strain. GAL4-UAS crosses were reared at 18°C and adult flies were transferred to 29°C to induce the transgenes before experiments. To assess the expression pattern of *egr* along the *Drosophila* midgut *egr-Gal4/CyO actGFP* (ref.18 Mabery & Schneider J Innate Immun. 2010) flies were crossed to *UAS-dsRed/TM3* (BDSC #6282) and *ry*^*506*^ *Dl-lacZ*^*05151*^*/TM3* (BDSC #11651) (ref. 30; Ranganavakulu et al Dev. Biol. 1996). Crosses were reared at 25°C and adult flies were transferred to 29°C for 2 days of aging before the experiment. The FlyFos line (VDRC# 318615) was used as an Egr-GFP protein trap (ref. 20; Sarov et al, eLife 2016).

### Oral infection assays

Performed as previously described (ref.11 Apidianakis et al PNAS 2009). Briefly, a single colony from the *P. aeruginosa* strain PA14 was grown at 37°C to OD_600nm_ = 3, corresponding to 5 × 10^9^ bacteria/mL. For immunohistochemistry, 20-30 newborn adult flies per genotype were aged for 5 days at 18°C and transferred for 5 d to 29°C to induce the Gal4 activity. Then flies were starved for 5 hours and added in groups of 10 per fly vial containing a cotton ball at the bottom impregnated with 5 ml of 0,5ml PA14, 1ml 20% sucrose and 3,5ml dH_2_O. For uninfected control 5ml of 1ml sucrose 20% and 4ml dH_2_O was used. Flies were incubated for 48 hours at 29°C.

### Survival assays

For individual *Drosophila* line testing, triplicates of ten 3-5 day old female flies for each extreme DGRP line were infected with the *P. aeruginosa* strain PA14, as described above. Dead and alive flies were recorded daily.

For the pooled strain survival assay, 5 female flies from each of the 11 highly mitotic DGRP strains containing the isogenized *UAS-srcGFP; Dl-Gal4* transgenes were mixed into a fly bottle with 5 female flies from each of the 11 lowly mitotic DGRP lines. The cohorts were infected in fly bottles containing 4 cotton balls at the bottom impregnated with 20 ml of 2ml PA14 culture (OD_600nm_=3), 4ml 20% sucrose and 14ml dH_2_O. Flies were incubated for 48 hours at 25°C. The number of dead and alive flies expressing GFP (highly mitotic) and not expressing GFP (lowly mitotic) was recorded daily with the use of the fluorescent Leica M165 FC stereoscope.

### Dissections and Immunohistochemistry

Performed as previously described (ref.11; Apidianakis et al PNAS 2009). Briefly, 15 midguts were dissected each time from each fly genotype in 1x PBS and fixed for 30 minutes with 4% formaldehyde (FA) at room temperature. Three quick rinses were performed with 1x PBS. Blocking with 1x PBS, 0.2% Triton-X, 0.5% BSA) for 20 minutes. Primary antibodies: rabbit-anti-pH3 (Millipore 1:4000), mouse-anti-Prospero (DSHB 1:100), mouse anti-β-gal (Promega 1:500), chicken-anti-GFP (Invitrogen 1:1000), rabbit anti-GFP (1:3000; Invitrogen) incubated overnight in the dark at 4°C. Midguts were washed 3 times for 10 minutes in 1x PBS, 0.2% Triton-X). Secondary antibodies against mouse, rabbit or chicken conjugated to Alexa fluor 488 and 555 (Invitrogen) were used at 1:1000. Samples were incubated in secondary antibody solution including DAPI (Sigma, 1:3000 of 10mg/ml stock) for 2 hours at room temperature, in the dark, with mild shaking. Midguts were washed 3 times. Mounting on glass microscope slides in 20μl of Vectashield (Vector), covered with glass coverslips and sealed with nail polish.

### Image acquisition and analysis

Stacks of optical sections were acquired using the Leica TCS SP2 DMIRE2 confocal microscope. Images to be compared were taken using the exact same settings. The numbers of pH3 cells were counted under the fluorescent microscope (Zeis Axioscope A.1) at 20x magnification along the whole midgut. For regional assessment of A1, A2, P1, P3 and P4 regions a standard frame of 300 × 350 µm per region per midgut was covered.

#### Midgut sizes

(length and width) were assessed from bright-field pictures acquired with a fluorescent Leica M165 FC stereomicroscope and their width and length was analyzed using Image J (https://imagej.nih.gov/ij/). Clicking on the Analyze, Set Scale option, using the length of the picture as 2048 pixels corresponding to 2.07mm. Using the segmented line option, a line was drawn manually, starting from the cardia and ending just before the Hindgut Proliferating Zone (HPZ). For the width measurement, straight lines indicating the width of posterior A1 and anterior P4 were manually drawn vertically to the gut length. The length of the lines was measured and thereby the gut’s dimensions, by clicking on the Analyze, Measure option.

#### Nucleus size

was calculated following analysis of confocal images using ImageJ. Confocal images of anterior (R2) and posterior (R5) midguts were captured at 40× magnification, zoom 1× and 1024×1024 format and produced as a maximum projection of 10-15 sections serial imaging. By clicking Analyze, Set Scale option, setting distance in pixels at 1024 and known distance at 375um, according to confocal photo properties. Multi-channeled images were subjected to the Split Channel option, to isolate the blue signal (DAPI staining). The images were subsequently converted into grey by selecting the Image option, Lookup Tables and Grey color option. Then, the adjustment of threshold was applied to produce 2-pixel intensities on the photo; black and white. A Binary version of the images was generated and the type of measurement was specified to the Analyze particles option: show outlines and infinity value set to either 1 or 2, to exclude the calculation of random speckles in the photo. Display Results, Clear Results, Summarize, Exclude Edges and Include Holes options were also selected. Measurements of area corresponding to each numbered nucleus were exported in Excel to calculate the mean area value. Merged nuclei were excluded manually to improve the efficiency of calculations.

### Statistical analysis

The Z-value of each strain was calculated by subtracting the average pH3 number of all 153 strains from the pH3 number of the strain, and dividing the subtraction by the standard deviation of the pH3 number of all 153 strains. It represents the number of standard deviations a particular DGRP line was found above or below the mean of all 153 lines.

To compare the means of two groups of values, namely, the pH3 positive numbers per midgut (n=30 midguts per genotype), the relative mRNA levels in “high” vs. “low” strains (n=11 strains × 3 biological replicates for each), CFUs (n=10 strains × 3 biological replicates for each), midgut dimensions (n=11 strains × 10 midguts for each), decoloration (n=11 strains x 3 biological replicates) and the nucleus area (n=11 strains × 3 midguts × 100 counts each) the two-tailed Student t-test was used. Data were visually inspected for normality of distribution with median being approximately in the middle of the 1^st^ and 3^rd^ quartile. A *p-*value < 0.05 is labeled as “*”, while *p*-values < 0.01 and < 0.001 are labeled as “**” and “***”, respectively. “ns” stands for not statistically significant. Error bars throughout represent standard deviation of the mean (STDEV).

Chi-square test was used to compare the number of total cell clusters of ≥5 cells per genotype between two genotypes sampling the same number of midguts (n=30 midguts) and expecting the same number of clusters per genotype. For d.f.=1 chi-square values were all >10,82 corresponding to significance with a *p*-value= 0.001, which is labeled as “***”. Error bars represent STDEV. Chi-square was also used to assess tumor incidence as being different among 3 genotypes (control, egrRNAi and egr oeverexpression) exhibiting 2, 1 and 7 tumors per 189, 167 and 144 midguts, respectively. For d.f.=2 the chi-square was >13,81 corresponding to significance with a *p*-value= 0.001.

For fly survival and decoloration curve assessment we applied the Kaplan-Meier method using the log-rank test (MedCalc statistical software).

Upon optimization each experiment was replicated independently for a total of 3 times. Results are presented taking into account all the replicates.

### RT-qPCR

For each of the 11 “high” vs 11 “low” extreme DGRP strains in both baseline and infected conditions the average of 3 biological replicates was used to assess the relative expression of regenerative inflammation genes (p values are given in Table 1). For Ras^V12^ expression experiment the average of 3 biological replicates x 3 technical replicates was used. RNA was extracted from 20 midguts per strain per condition per biological replicate using Qiazol. 800 ng of total RNA were used to synthesize the cDNA using Promega RQ1 RNase-Free DNase Kit according to the manufacturers’ protocol. Reverse transcription was performed using 145,4ng of the total DNAse treated RNA by using the TaKaRa Prime ScriptTM RT Master Mix Kit. qPCR amplification was performed using gene specific primers with the following amplification program: 95°C for 30 seconds (initial denaturation), 40 cycles of 95°C for 10 seconds (denaturation), 60°C for 30 seconds (annealing), 65°C for 30 seconds (extension) and 65°C for 1 minute (final extension). Primer sequences for each gene are shown in Table 3. The expression of the genes of interest was normalized to the expression levels of two reference genes, *rpl32* and *gapdh1* using the 2^-ΔΔCt^ method. Data were analyzed using the Bio-rad CFX Manager 3.1 program.

**Table 3:**
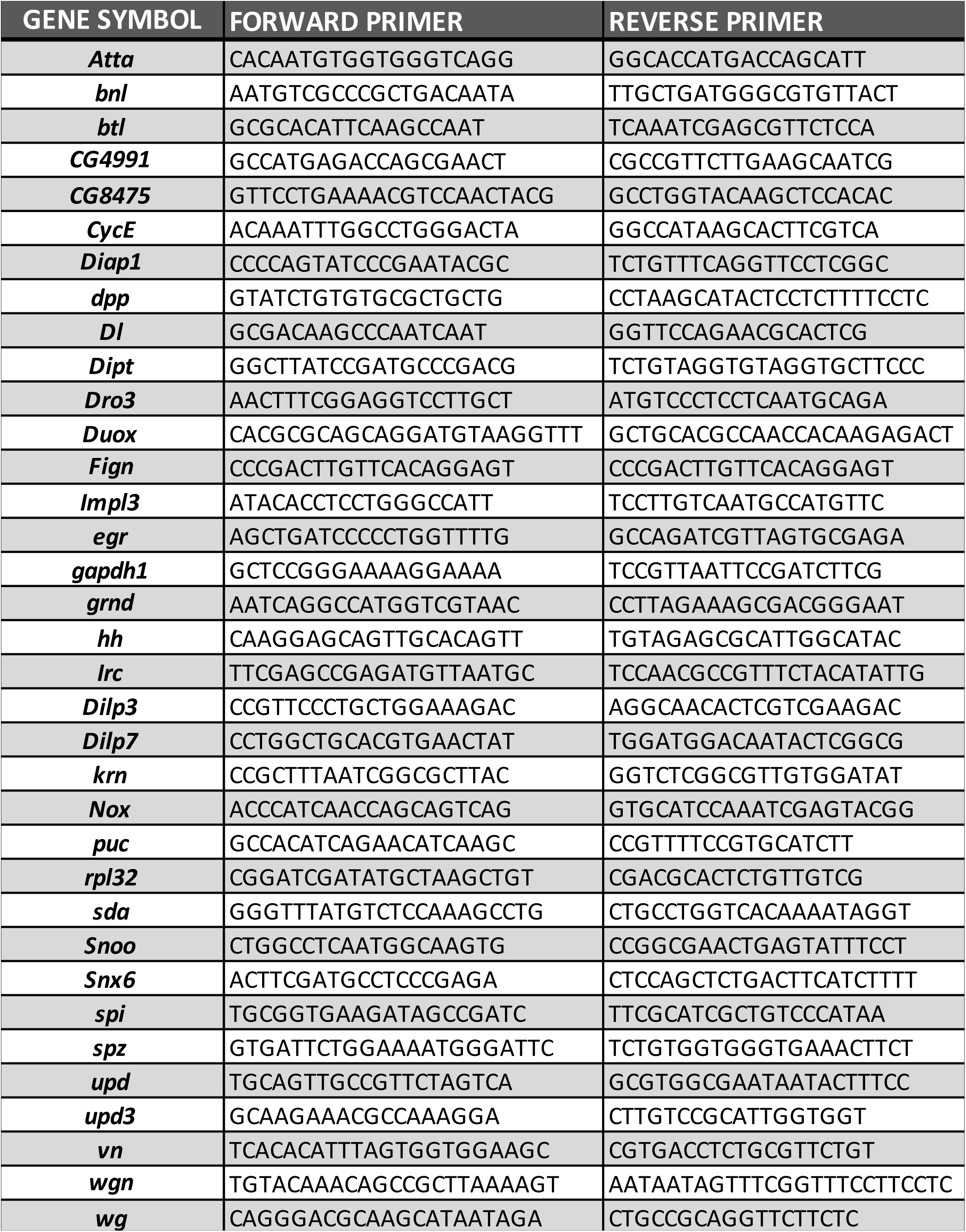
List of primers used in RT-qPCR experiments.

### Dysplastic cell cluster enumeration

*w; UAS-srcGFP; Dl-Gal4* flies were backcrossed to the 22 extreme DGRP lines for 6 generations by selecting GFP+ larvae in each generation to obtain the genetic background of the original DGRP line, while having GFP marked ISCs. 5-7 days old flies from each strain were orally infected with the *P. aeruginosa* for 24 hours at 25°C as described above and transferred into 50ml falcon tubes bearing 12 1.2mm holes on the lid (for access to food) and 32 0.5mm holes on the tube surface (for aeration), using flame heated 18g × 40mm and 25g × 16mm needles to pierce the tube lid and surface, respectively. A Whatman disc (23mm) (Sigma Aldrich) impregnated with 270μl of a solution composed of 1mM DAPT (Sigma Aldrich) dissolved in 30% yeast paste was placed on the outside of the tube lid and stabilized with parafilm. Flies were treated with DAPT for 4 days at 25°C and flipped every day into clean falcons with freshly prepared drug.

### Methylene blue – EC coloration

5-7 day old adult females were fed on 0,5% Methylene Blue (Sigma) dissolved in 85% heat-killed yeast paste for 5 hours at 25°C secondary to 5 hours starvation. Flies were then subjected to either bacterial infection or 4% sucrose feeding for 2 days. Then, flies were fed on 4% sucrose and recorded everyday according to their color status (blue versus non-blue abdomen) until complete decoloration of their guts was observed.

### Bacterial load

*P. aeruginosa* (PA14 strain) colony forming units (CFUs) per fly strain were determined following 2 days of infection at 25°C. Flies were externally sterilized by brief dipping into pure ethanol, dried, and placed into 2ml eppendorf tubes containing 200μl LB and a stainless steel bead of 5mm diameter (Qiagen). Flies were homogenized using the TissueLyser II (Qiagen) at 50Hz for 5 minutes. LB was then added into the tubes containing the tissue lysate to reach the volume of 1000μl. Serial dilutions of the lysate obtained from three flies were plated onto LB agar plates selective for PA14 containing 100μl/ml Rifampicin (Sigma) and incubated overnight at 37°C. In total, bacterial colonies from three replicates per DGRP extreme line were counted.

### Germ-free flies

Flies were transferred in empty bottles covered with a fruit juice agar plate (35mm × 10mm). The fruit juice agar plate was prepared following boiling of 2% agar dissolved in fruit juice and supplemented with Tegasept and Propionic acid to a final concentration of 0,56% and 0,37%, respectively. Once the mixture was solidified, 0,2ml of yeast paste (66% dry yeast dissolved in ddH2O) was transferred in the middle of each petri dish. Flies were conditioned by feeding on fruit juice agar plates for a day before transferred into clean bottles with freshly prepared fruit juice agar plates on the top. After a 15 hour incubation at 25°C the eggs were collected into a mesh basket using a brush. Each basket was placed in a beaker containing 20ml of 50% bleach for a maximum of 2 minutes or until ∼80% of dorsal appendages were dissolved due to the removal of the chorion layer. Bleached eggs were then washed with sterile ddH2O under the microbiological hood and transferred into bottles containing sterile fly food and maintained at 25°C. Once the offspring began to emerge, it was transferred into bottles with sterile food. Lysates obtained from the emerged flies were plated onto LB media and incubated at 37°C overnight to ensure that they were germ-free.

### *Drosophila* Aging experiments

Flies were maintained at 25°C on our standard yeast-cornmeal-sucrose food. *w/ UAS-srcGFP; Dl-Gal4;* and *OreR/UAS-srcGFP; Dl-Gal4* flies were produced by crossing *w; UAS-srcGFP/CyO; Dl-Gal4/TM6C* to *w*^*1118*^ and Oregon R. DGRP lines #28194 and #28217 with GFP marked ISCs were produced by backcrossing *UAS-srcGFP/+;Dl-Gal4/+* flies to the original DGRP lines for 6 generations and selecting GFP+ larvae in each generation. Following mating for 4 days males and females were kept separated, 20 flies per vial and flipped on fresh food every two days for the first 20 days and then every day from day 20 to 42 day. ISC/EE clusters of 5, 6, 7 and ≥8 cells and anti-phospho-histone H3 (pH3) reactivity were measured from female flies stained with anti-GFP and anti-Prospero or anti-pH3 at 4, 30 and 42 days. For tumor detection, male flies of one cohort per genotype were dissected at 42 days of age stained with anti-GFP, anti-Prospero and DAPI for tumor assessment. Tumors were defined as masses of >100 cells that are GFP+ and/or Prospero+ cells and have smaller nuclei than mature ECs.

### Unique material availability statement

All unique materials used are commercially available or readily available by the authors.

